# Acute stress causes sex-dependent changes to ventral subiculum synapses, circuitry, and anxiety-like behavior

**DOI:** 10.1101/2024.08.02.606264

**Authors:** Carley N Miller, Yuan Li, Kevin T Beier, Jason Aoto

**Affiliations:** Department of Pharmacology, University of Colorado Anschutz Medical Campus, Aurora, CO 80045, USA; Department of Physiology and Biophysics, University of California, Irvine, CA, USA 92697; Department of Neurobiology and Behavior, University of California, Irvine, CA, USA 92697; Department of Biomedical Engineering, University of California, Irvine, CA, USA 92697; Department of Pharmaceutical Sciences, University of California, Irvine, CA, USA 92697

## Abstract

Experiencing a single severe stressor is sufficient to drive sexually dimorphic psychiatric disease development. The ventral subiculum (vSUB) emerges as a site where stress may induce sexually dimorphic adaptations due to its sex-specific organization and pivotal role in stress integration. Using a 1-hr acute restraint stress model, we uncover that stress causes a net decrease in vSUB activity in females that is potent, long-lasting, and driven by adrenergic receptor signaling. By contrast, males exhibit a net increase in vSUB activity that is transient and driven by corticosterone signaling. We further identified sex-dependent changes in vSUB output to the bed nucleus of the stria terminalis and in anxiety-like behavior in response to stress. These findings reveal striking changes in psychiatric disease-relevant brain regions and behavior following stress with sex-, cell-type, and synapse-specificity that contribute to our understanding of sex-dependent adaptations that may shape stress-related psychiatric disease risk.

**Highlights:**

- vSUB BS cells are uniquely stress sensitive
- Stress causes sex-dependent changes to BS cell E/I balance
- Stress causes sex-dependent changes to vSUB activity to aBNST *in vivo*
- Stress causes anxiety-like behavior in females, but not males

## Introduction

Stress exposure is a prominent risk factor for developing psychiatric disease and females experience stress-related psychiatric diseases at substantially greater rates than males^1^. Further, stress-related psychiatric diseases exhibit profound sexual dimorphism, making it critical to understand the mechanisms that shape sex-dependent vulnerability. Chronic stress is well understood for its contribution to psychiatric disease development in both sexes^2^ and causes synaptic adaptations within the ventral hippocampus (vHipp) in preclinical and clinical models.^3,4^ Interestingly, severe acute stress is a critical risk factor for psychiatric disease development disproportionately in females, intimating that there may be sex differences in how acute stress is encoded at synapses relevant to psychiatric disease pathogenesis.^5–8^ However, the cellular mechanisms by which acute stress induces vulnerability to psychiatric disease remain obscure, and even less is understood regarding how acute stress impacts vHipp synapses while considering sex as a biological variable.

Excitatory/inhibitory (E/I) imbalance within vHipp has been firmly established in chronic stress and psychiatric disease models in both sexes,^9^ revealing altered synaptic plasticity at excitatory synapses and in the excitability of parvalbumin (PV) expressing inhibitory interneurons.^10–13^ The ventral subiculum (vSUB) subregion of vHipp emerges as a promising site of sex-specific disease pathogenesis due to its basal sexually dimorphic microcircuit organization and its regulation of stress integration and anxiety-like behavior via projections to the anterior bed nucleus of the stria terminalis (aBNST).^14–17^ Although vSUB is sexually dimorphic and resides at the intersection of stress regulation and anxiety-like behavior, vSUB is grossly understudied.

Distinct from other vHipp subregions, vSUB primarily consists of pyramidal excitatory cells that are classified based on their firing pattern as burst (BS) or regular spiking (RS). RS and BS cells are believed to be functionally nonoverlapping, especially in psychiatric disease pathogenesis.^18–22^ These cells express receptors targeted by the main stress effector pathways, the sympathetic nervous system (SNS) and hypothalamus-pituitary-adrenal gland axis (HPA).^23–26^ While SNS and HPA activity are vital to respond appropriately to environmental stressors, dysfunction of either system can contribute to the development of psychiatric disease.^27,28^ Together, the inherent sexual dimorphism and stress responsivity of vSUB make this subregion well-positioned as a site of sex-dependent stress integration and stress disorder pathogenesis; however, no studies have systematically examined vSUB synapses after acute stress with cell-type and sex specificity.

Here, using multidisciplinary approaches, we systematically examined the impact of 1-hr acute restraint stress (ARS) on the functional properties of vSUB synapses in *ex vivo* slices, vSUB activity *in vivo*, and anxiety-like behavior in male and female mice. ARS induced sex-, cell-type- and synapse-specific adaptations in the E/I balance of vSUB principal cells that resulted in net inhibition of BS cells in females and net excitation of BS cells in males.

Importantly, these sex-specific functional adaptations in response to ARS were mechanistically distinct as adrenergic receptor signaling was necessary in females while corticosterone signaling was required in males. Further, the synaptic adaptations to ARS in females were immediate and long-lasting while in males, the synaptic adaptations were delayed and transient. Moreover, our electrophysiological and *in vivo* calcium imaging identified sex-specific ARS-induced changes in vSUB output to the aBNST. Finally, ARS produced anxiety-like behavior in female, but not male mice, and provides the framework to interpret the *ex vivo* and *in vivo* synaptic findings we find in parallel. Our results endorse that acute stress integration is sex-dependent in psychiatric disease-relevant brain regions and represent initial steps to understand the mechanisms by which sex differences in stress-induced psychiatric disease prevalence, treatment responsivity, and symptomology may occur.

## Results

### Stress impairs CA1-vSUB-BS excitatory synapses in females

We first interrogated the strength of vCA1-BS and vCA1-RS excitatory synapses 24-hrs after a 1-hr ARS paradigm. vCA1 provides the majority glutamatergic input onto vSUB RS and BS principal neurons, and dysfunction at vCA1-vSUB synapses may cause an E/I imbalance and drive a pathologic state.^9^ We monitored electrically-evoked excitatory postsynaptic currents (EPSCs) from electrophysiologically identified postsynaptic RS and BS neurons in *ex vivo* acute vHipp slices (Figures 1A-B). To selectively isolate vCA1 afferents to vSUB, we placed our extracellular stimulation electrode in the alveus/stratum oriens border of vCA1 (Figures 1B). In females, ARS reduced vCA1-BS synaptic strength by over 50% according to their input-output relationship (Figure 1d). We next measured strontium-evoked asynchronous EPSCs (aEPSCs) and EPSC paired-pulse ratios (PPRs) to identify the synaptic locus of this stress-induced decrease in excitatory transmission. aEPSC amplitudes are commonly believed to reflect postsynaptic strength^29^ while PPRs are an indirect measure of presynaptic release. In females, ARS reduced aEPSC amplitudes at vCA1-BS synapses without altering the PPR, indicating that ARS selectively decreased postsynaptic strength at these synapses (Figures 1E-F). At vCA1-RS synapses, ARS induced a modest, but statistically significant, decrease in postsynaptic strength (Figures 1G-J). Together, these stress-induced changes in vCA1-vSUB synapses manifest postsynaptically and were greater in BS cells, consistent with the current notion that BS cells are more susceptible to stress-induced perturbations.^25,30,31^

**Figure 1.**
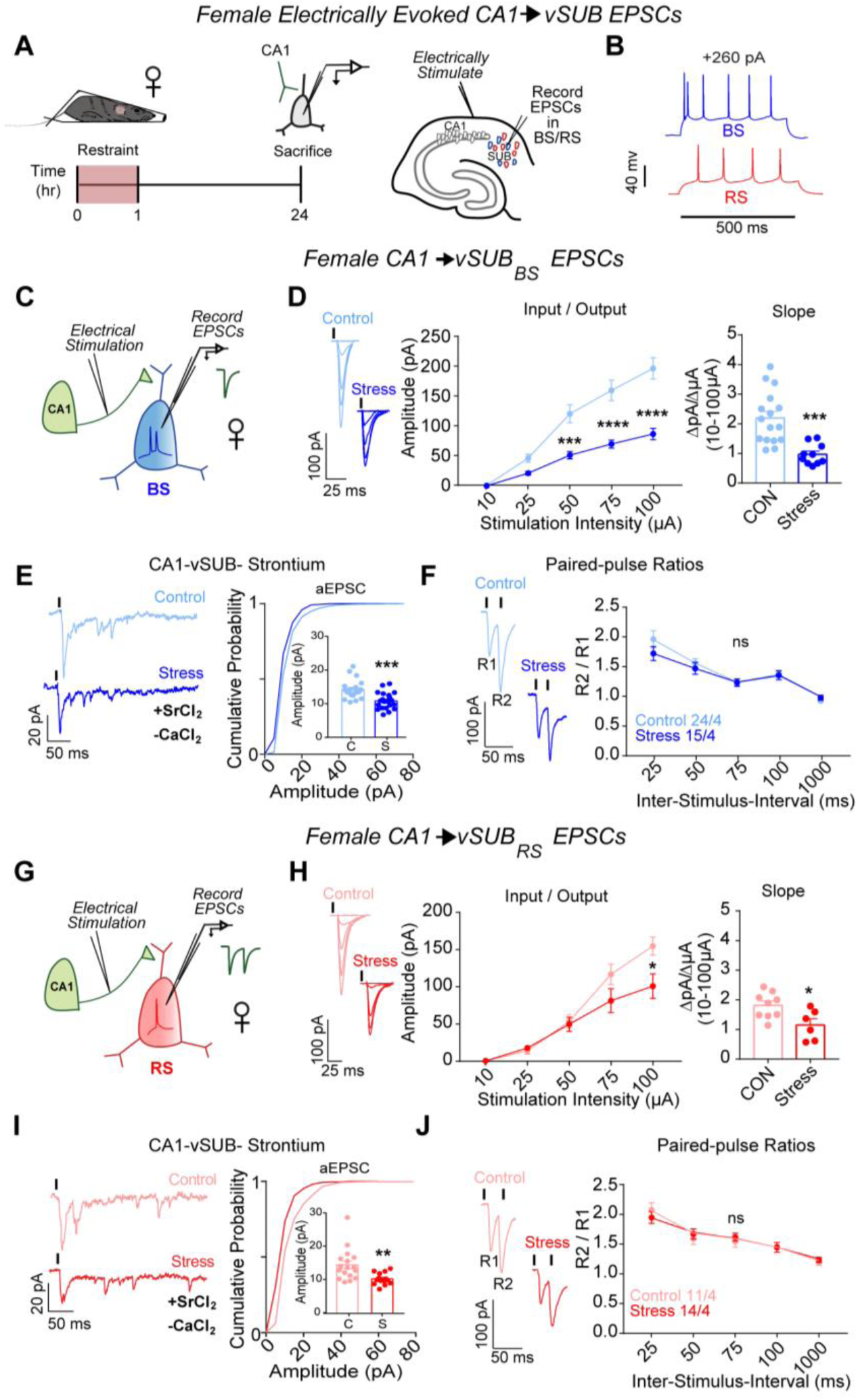
ARS weakens CA1-vSUB basal excitatory synaptic strength in females. (**A**) Electrically evoked EPSCs from CA1 were recorded in vSUB neurons 24-hr after stress. (**B**) Representative traces of action potential firing patterns in BS (blue) or RS (red) cells. (**C**,**G**) Electrically stimulated EPSCs were recorded in BS or RS cells. (**D**) Input-output representative traces (left), summary graph (middle), and slope (right) for EPSCs recorded in BS (10 μA, *P*>0.9999; 25 μA, *P*=0.5756; 50 μA, \*\*\**P*=0.0007; 75 μA, \*\*\*\**P*<0.0001; 100 μA, *\*\*\*\*P*<0.0001; slope, \*\*\**P*=0.0003; control N=16/4, stress N=10/4 ) or (**H**) RS cells ( 10 μA, *P>*0.9999; 25 μA, *P*=0.9996; 50 μA, *P*=0.9996, 75 μA, *P*=0.0828; 100 μA, \**P*=0.0022; slope, *P*=0.0189; control n=9/3, stress n=6/4). (**E,I**) Representative traces of strontium-mediated aEPSCs after electrical stimulation (left) and aEPSC amplitude (right) for BS (**E**; \*\*\**P*= 0.0002; control n=20/3, stress n=21/3) or RS cells (**I**; \**P*=0.043; control n=17/3, stress n=14/3). (**F**,**J**) Representative PPR trace (50ms) (left) and PPR measurements (right) from BS (**F**; 25-ms ISI, *P*=0.2461; 50-ms ISI, *P*= 0.9573; 75-ms ISI, *P*>0.9999; 100-ms ISI, *P*>0.9999; 1000-ms ISI, *P*=0.9959; control n=24/4, stress n=15/3) or RS cells (**J**; 25-ms ISI, *P*=0.8008; 50-ms ISI, *P*=0.9785; 75-ms ISI, *P*=0.9827; 100-ms ISI, *P*>0.9999; 1000-ms ISI, *P*=0.9990; control n=11/4, stress n=14/4). Data are represented as mean +/- SEM; means were calculated from the total number of cells. Numbers in the legend represent the numbers of cells/animals. Statistical significance was determined by a 2-way ANOVA or unpaired t-test. See also Figure S1.

The prominent ARS-dependent reduction in excitatory synaptic transmission at vCA1-BS synapses in females motivated us to examine whether similar stress-induced synaptic adaptations occur in male mice. In contrast to females, ARS did not impact vCA1-vSUB synaptic strength, postsynaptic strength in BS or RS cells of males, or presynaptic release from vCA1 (Figures 2A-J), consistent with other models of acute stress.^25,32^ These data reveal that ARS causes sexually dimorphic changes to basal excitatory strength in vSUB and prompted us to examine whether ARS also alters activity-dependent plasticity.

**Figure 2.**
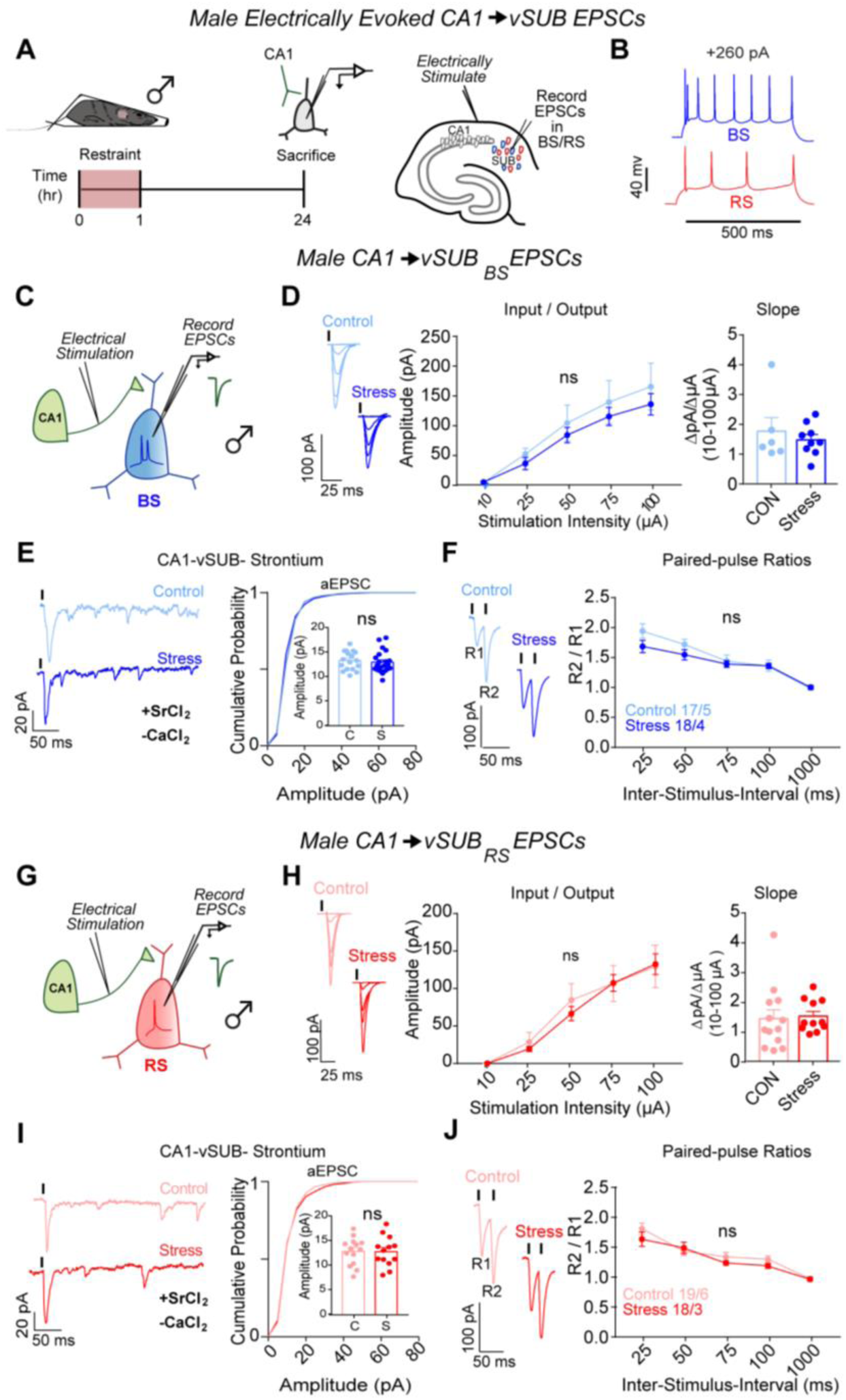
ARS does not alter CA1-vSUB basal excitatory synaptic strength in males. (**A**) Electrically evoked EPSCs from CA1 were recorded in vSUB neurons 24hr after stress. (**B)** Representative traces of firing patterns in BS or RS cells. (**C**,**G**) Electrically stimulated EPSCs were recorded in BS or RS cells. (**D**) Input-output representative traces (left), summary graph (middle), and slope (right) for EPSCs recorded in BS (**D**; 10 μA, *P*>0.9999; 25 μA, *P*=0.9933; 50 μA, *P*=0.9887; 75 μA, *P=*0.9863; 100 μA, *P=*0.9701; slope, *P*=0.7043; control n=6/3, stress n=8/3) or (**H**) RS cells (10 μA, *P>*0.9999; 25 μA, *P*=0.9976; 50 μA, *P*=0.9484, 75 μA, *P>*0.9999; 100 μA, *P>*0.9999; slope, *P*=0.8081; control n=13/3, stress n=11/4). (**E**,**I**) Representative traces of strontium-mediated aEPSCs after electrical stimulation (left) and aEPSC amplitude (right) for BS (**E**; *P*= 0.6312; control n=23/3, stress n=14/3) or RS cells (**I**; *P*=0.9679; control n=19/3, stress n=21/3). (**F**,**J**) Representative PPR trace (50ms) (left) and PPR measurements (right) from BS (**F**; 25-ms ISI, *P=*0.1233; 50-ms ISI, *P*= 0.5243; 75-ms ISI, *P=*0.9957; 100-ms ISI, *P*>0.9999; 1000-ms ISI, *P>*0.9999; control n=17/5, stress n=18/4) or RS cells (**J**; 25-ms ISI, *P=*0.3803; 50-ms ISI, *P*= 0.9821; 75-ms ISI, *P=*0.8702; 100-ms ISI, *P=*0.8262; 1000-ms ISI, *P>*0.9999; control n=19/6, stress n=18/4). Data are represented as mean +/- SEM; means were calculated from the total number of cells. Numbers in the legend represent the numbers of cells/animals. Statistical significance was determined by a 2-way ANOVA or unpaired t-test. See also Figure S2.

### Stress impairs LTP at vCA1-BS synapses in females

Hippocampal plasticity occurs in response to major stressors^33–35^ and the dysfunction of plasticity is a hallmark of psychiatric diseases. Long-term potentiation (LTP) in vSUB manifests pre- or post-synaptically depending on the firing pattern of the postsynaptic cell. CA1-BS LTP occurs presynaptically due to increased presynaptic release probability, while CA1-RS LTP manifests postsynaptically through classical NMDA-receptor-dependent mechanisms^14^. Using an established induction protocol,^21,36^ we assayed LTP at vCA1-BS and vCA1-RS synapses in females after ARS. ARS impaired presynaptic LTP in BS cells (Figures S1A-D) without altering postsynaptic LTP in RS cells (Figures 1E-F), reinforcing a stress-sensitive characterization of BS cells. By contrast, in males, LTP was intact at vCA1-BS and vCA1-RS synapses (Figures 2A-F). Our results indicate that ARS impairs vCA1-BS synapses in females through two distinct routes: 1. A reduction of basal postsynaptic strength and 2. Rendering this synapse incapable of activity-dependent presynaptic LTP. Together, ARS causes sex-specific excitatory synaptic adaptations such that excitatory basal synaptic transmission and activity-dependent plasticity are remarkably impaired after ARS in the vSUB of only females, and vSUB BS synapses are dominantly impacted.

### Stress causes sexually dimorphic adaptations to vSUB PV-BS inhibition

We next probed whether ARS drives changes in PV inhibition of BS cells to compensate for or exacerbate the robust cell-type-specific impairment of basal excitatory transmission and LTP on BS cells in females. PV interneurons exhibit sex-specific connectivity in vSUB, are more stress-sensitive than other hippocampal interneurons, and exert major inhibitory control over vSUB pyramidal cells.^17, 37–41^ Despite this, how stressors impact PV synaptic inhibition in any brain region remains unexamined. To isolate PV inhibition, we injected a Cre-dependent channelrhodopsin variant (AAV-DIO-ChIEF) into the vSUB of P21 PV-Cre mice and assessed light-evoked PV inhibitory postsynaptic currents (IPSCs) from RS and BS cells in acute slices from adult mice after ARS (Figures 3A-B). In females, ARS significantly increased PV-BS inhibition by enhancing postsynaptic strength without altering presynaptic release (Figures 3C-F). By contrast, ARS did not significantly change PV-RS inhibition (Figures 3H-J). In males, ARS surprisingly reduced the postsynaptic strength of inhibitory PV-BS synapses without altering presynaptic release (Figures 4 A-F). Thus, although ARS induced distinct sex-specific changes in the inhibitory strength of PV-BS synapses, both phenotypes share a postsynaptic locus. Similar to females, ARS did not impact PV-RS inhibition in males (Figures 4G-J). Together, ARS increases the strength of PV-BS synapses and reduces vCA1-BS excitatory synaptic strength, resulting in an overall net reduction in vSUB-BS activity in females. By contrast, ARS reduces PV-BS inhibition without changes in vCA1-BS excitatory strength in males which may shift vSUB-BS cells to a more active state and provide a synaptic explanation for why the general activity of vSUB is increased after ARS in males.^24,42^

**Figure 3.**
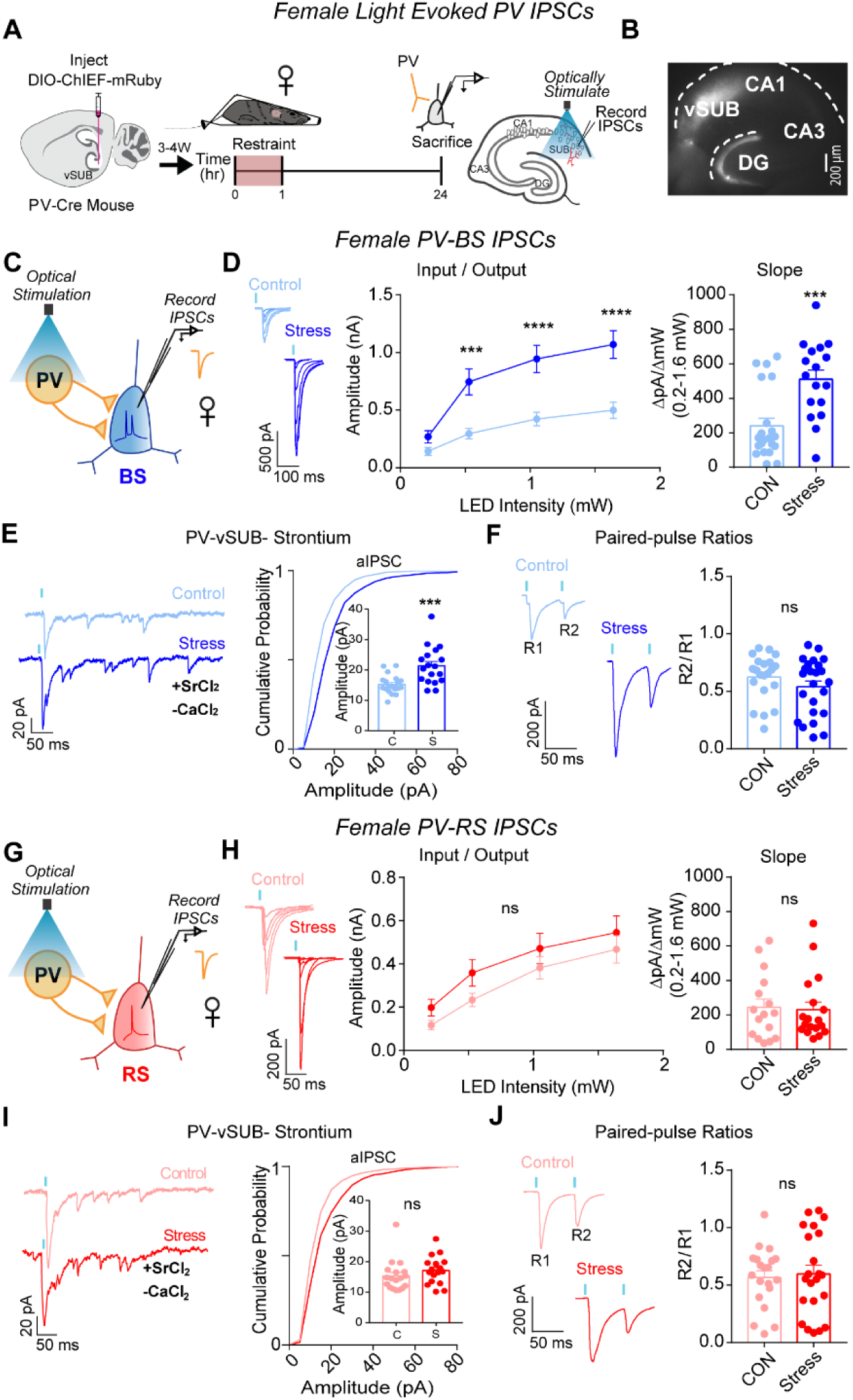
ARS strengthens PV-vSUB-BS inhibition via increased postsynaptic strength in females. (**A**) A Cre-dependent ChIEF AAV was injected into vSUB of PV-Cre female mice and optogenetically evoked IPSCs from PV interneurons were recorded from BS or RS cells 24-hr after 1-hr of ARS. (**B**) Example image of ChIEF-mRuby expression in the vSUB. (**C,G**) Optogenetically stimulated EPSCs were recorded in BS or RS cells after stress. (**D,H**) Input-output representative traces (left), summary graph (middle), and slope (right) for IPSCs recorded in BS (**D**; 0.213 pA, *P*=0.7032; 0.518 pA, \*\*\**P*=0.0004; 1.050 pA, \*\*\*\**P<*0.0001; 1.640 pA, \*\*\*\**P*<0.0001; slope, \*\*\**P*=0.0004; control n=21/3, stress n=17/3) or RS cells (**H**; 0.213 pA, *P*=0.7696; 0.518 pA, *P*=0.3993; 1.050 pA, *P*=0.7114; 1.640 pA, *P*=0.7976; slope, *P*=0.8366; control n=16/3, stress n=18/3). (**E,I**) Representative traces of strontium-mediated aIPSCs after optogenetic stimulation (left) and aIPSC amplitude (right) for BS (**E**; \*\*\**P*=0.0004; control n=20/3, stress n=18/3) or RS cells (**I**; *P*=0.2681; control n=18/3, stress n=17/3). (**F**,**J**) Representative PPR traces (50ms) (left) and PPR measurements (right) from BS (**F**; *P*=0.2175; control n= 21/3, stress n= 24/3) or RS cells (**J** *P*=0.7608; control 20/3, stress n=22/3). Data are represented as mean +/- SEM; means were calculated from the total number of cells. Numbers in the legend represent the numbers of cells/animals. Statistical significance was determined by a 2-way ANOVA or unpaired t-test. See also Figures S3, S5, S7.

**Figure 4.**
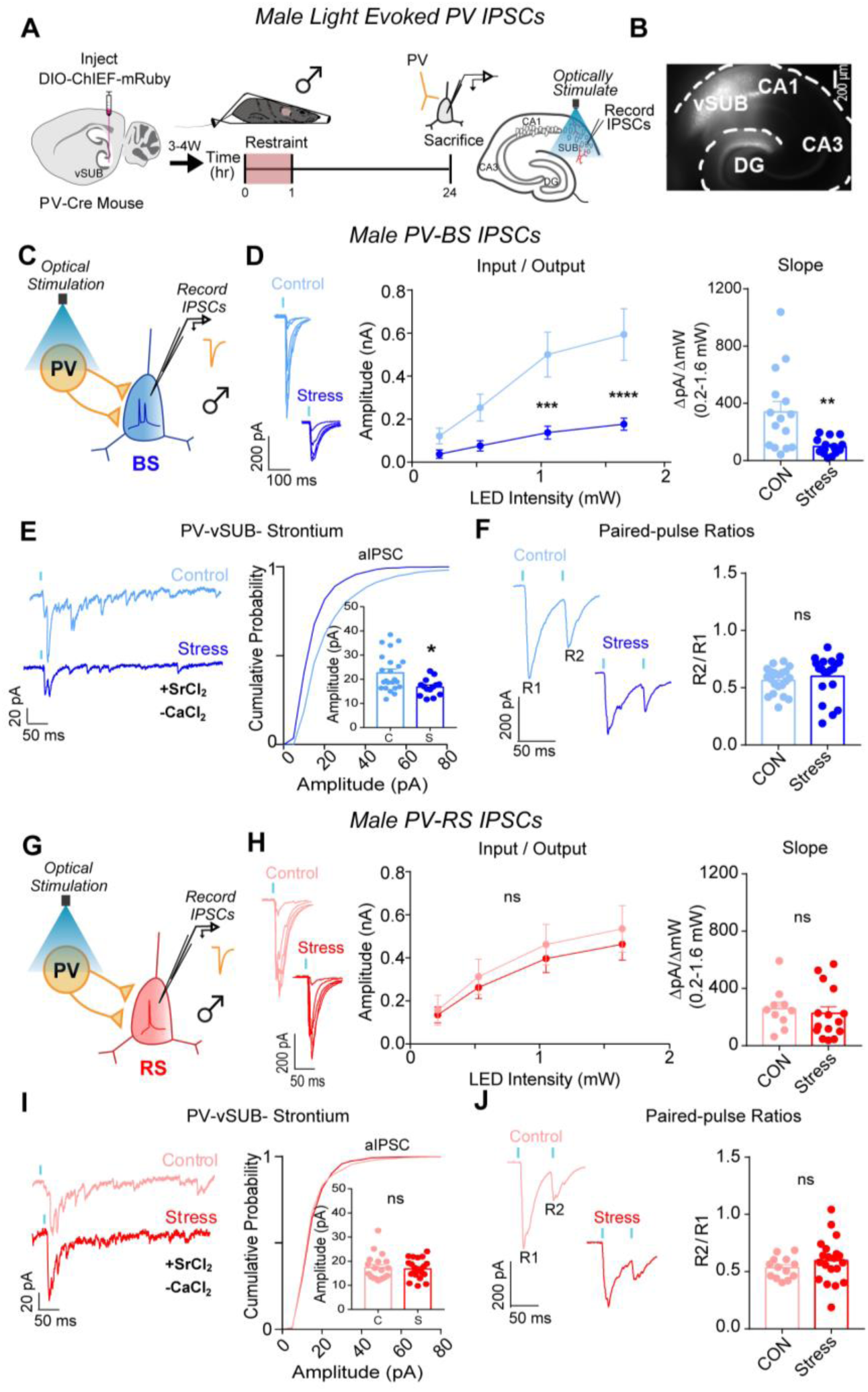
ARS weakens PV-vSUB-BS inhibition via decreased postsynaptic strength in males. (**A**) A Cre-dependent ChIEF AAV was injected into vSUB of PV-Cre female mice and optogenetically evoked IPSCs from PV interneurons were recorded from BS or RS cells 24 hr after 1 hr of restraint stress. (**B**) Example image of ChIEF-mRuby expression in the vSUB. (**C,G**) Optogenetically stimulated EPSCs were recorded in BS or RS cells after stress. (**D,H**) Input-output representative traces (left), summary graph (middle), and slope (right) for IPSCs recorded in BS (**D**; 0.213 pA, \*\*\*\**P<*0.0001; 0.518 pA, \*\*\*\**P<*0.0001; 1.050 pA, \*\**P*=0.0030; 1.640 pA, \*\*\*\**P*<0.0001; slope, \*\**P*=0.0029; control n=15/3, stress n=15/3) or RS cells (**H**; 0.213 pA, *P*=0.9992; 0.518 pA, *P*=0.9813; 1.050 pA, *P*=0.9491; 1.640 pA, *P*=0.9289; slope, *P*=0.6343; control n=10/3, stress n=15/3). (**E,I**) Representative traces of strontium-mediated aIPSCs after optogenetic stimulation (left) and aIPSC amplitude (right) for BS (**E**; \*\**P*=0.0109; control n=24/3, stress n=16/3) or RS cells (**I**; *P*=0.7594; control n=19/3, stress n=21/3). (**F**,**J**) Representative PPR traces (50ms) (left) and PPR measurements (right) from BS (**F**; *P*=0.4339; control n= 21/3, stress n= 24/3) or RS cells (**J**; *P*=0.2890; control 13/3, stress n=19/3). Data are represented as mean +/- SEM; means were calculated from the total number of cells. Numbers in the legend represent the numbers of cells/animals. Statistical significance was determined by a 2-way ANOVA or unpaired t-test. See also Figures S4, S6, S7.

Stress can alter the activity of brain regions by modulating the intrinsic excitability of cells,^43–45^ however, these studies have primarily examined males. We measured action potential (AP) frequency and rheobase from BS, RS, and PV neurons in vSUB in mice from both sexes. In females, ARS did not alter the intrinsic excitability of vSUB BS, RS, or PV cells (Figures S3A-C). Similarly, in males, ARS did not alter the intrinsic excitability in BS or RS cells (Figures S4A-B). However, ARS reduced AP frequency and increased the rheobase of PV cells in males (Figures S4C). While these changes likely impair the sustained firing of male PV interneurons, they are unlikely to directly account for the reduction in the amplitudes of single light-evoked IPSCs at PV-BS synapses because our data indicate that this phenotype is driven by a reduction in postsynaptic strength.

### Stress causes lasting changes to PV-BS inhibition and vSUB cellular identity in females

The diametrically opposing adaptations to PV-BS inhibition 24-hrs after ARS in males and females led us to ask whether these sex-specific effects reflect transient or long-lasting changes to vSUB circuitry. To our knowledge, a time course of the susceptibility and sensitivity of vSUB BS cells to ARS is untested. In females immediately after ARS, the postsynaptic strength of PV-BS synapses was enhanced and closely mirrored what we observed at 24-hr post-stress (Figures S5A-D). Further, there were no changes in PV-RS inhibition (Figures S5E-G). The immediacy and magnitude of the effects of stress on PV-BS were strikingly potent in females, further underscoring these synapses as highly stress-sensitive. In comparison, the reduction in PV-BS inhibition observed 24-hrs post-stress in males (Figures 4A-F) was not present immediately after ARS (Figures S6A-F). Together, female PV-BS synapses more rapidly adapt to ARS than males in the acute window following ARS.

We next examined whether the ARS-induced adaptation at female PV-BS synapses persisted 1-week post-stress (Figures S7A-B). Curiously in females, 1-week after ARS, ∼97% (31 of 32 vSUB cells) exhibited burst-spiking properties (Figure S7c). This was unexpected because it is well-accepted that vSUB BS and RS neurons exist in roughly equal proportions.^46^ Using the same experimental approach in age- and littermate-matched control females, BS and RS cells were equally prevalent. This strongly suggests that during the 1-week after ARS, female vSUB RS cells adopted a burst-spiking profile (Figure S7C). This conversion was sex-specific and not observed in males (Figure S7G). Conversion from RS to BS cells in other brain regions has been implicated in the pathophysiology of anxiety^47,48^ and depression^49^ and has been observed after long-term social defeat in the subiculum of male mice.^22,50^ Thus, this ARS-induced long-term increase in bursting activity of female vSUB neurons reflects stress-specific adaptations and may contribute to the susceptibility of stress-related disorders.

1-week after ARS in females, the strength of PV-BS synapses was increased due to enhanced postsynaptic strength, consistent with the immediate and 24-hrs timepoints (Supplementary Fig. 7d-f). By contrast in males, 1-week after stress there were no significant differences in PV-BS or PV-RS inhibition (Figures S7h-n). In sum, the impact of ARS on PV-BS synapses is incredibly potent and lasting in females. Divergently in males, ARS induces a brief window of plasticity at PV-BS synapses. These data demonstrate that ARS produces profound sex-dependent changes in PV-BS inhibitory synaptic phenotypes, the time course of these phenotypes, and the cellular identity of vSUB pyramidal cells. Our findings emphasize the vSUB as a site for further examination to understand sex differences in stress-induced pathophysiology.

### vSUB-BS cells disproportionately project to aBNST

The aBNST is a highly sexually dimorphic nucleus that controls stress and anxiety states.^51,52^ The vSUB-aBNST circuit is known to be critical for stress hormone (i.e. corticosterone) release and anxiety-like behavior, but its cell type-specific connectivity remains undefined. To determine the cellular identity of aBNST-projecting cells within the vSUB, we stereotactically injected mRuby expressing retrograde AAV2 into the aBNST of P21 mice. To gain cell-type resolution, we electrophysiologically defined mRuby-positive vSUB neurons. Despite the vSUB containing RS and BS in equal proportions,^19,46^ we found that ∼80% of aBNST-projecting neurons in the vSUB were BS in both sexes (Figures 5A-E). Given this notable BS cell over-representation, we assessed whether there were differences in intrinsic excitability between RS and BS cells that project to aBNST that may compensate for this bias. We found no differences in RS or BS cell intrinsic excitability in either sex (Figures 5F-G). Our data show that BS neurons are uniquely stress-sensitive and convey most information from the vSUB to aBNST in both sexes.

**Figure 5.**
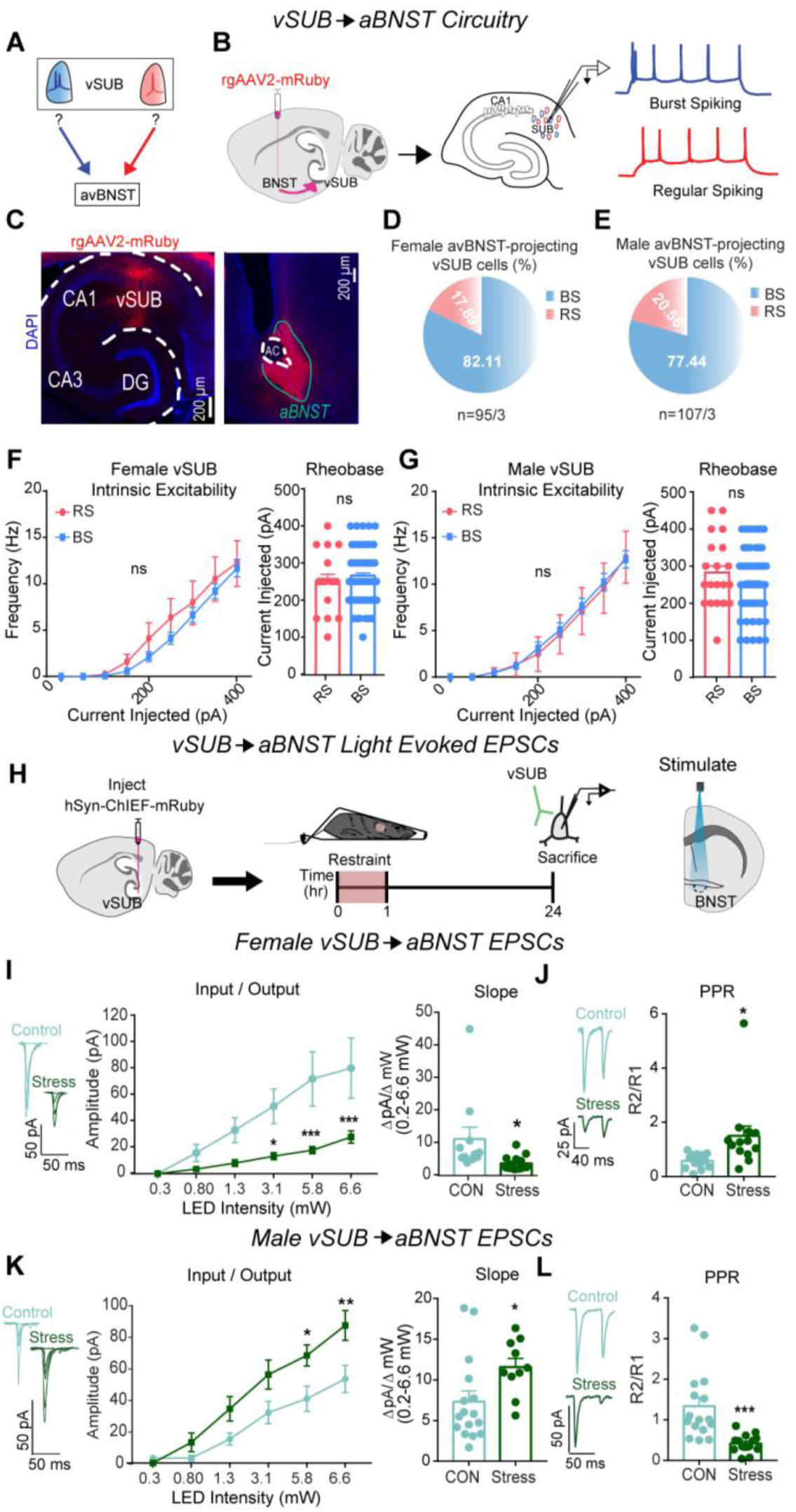
vSUB output to the aBNST is primarily from BS cells and stress causes sex dependent changes in vSUB-aBNST synaptic strength. (**A**) Diagram depicting the unknown identity of vSUB cells that project to the aBNST. (**B**) Diagram of injection of rgAAV2-mRuby in the aBNST and subsequent patching of fluorescently labeled cells in the vSUB to identify cells as BS or RS. (**C**) Representative images of the mRuby expression in the vSUB and injection site in the aBNST. (**D,E**) Cellular identity of aBNST projecting cells in the vSUB of female (n=95/3) or male (n=107/3) mice. (**F-G**) Intrinsic excitability (left) and rheobase (right) of aBNST projecting cells in the vSUB in female (**f**; *P*=0.3398; rheobase, P=0.6624; BS=64/3 RS=16/3) or male (**G**; *P*=0.5432; rheobase, *P*= 0.1793; BS=72/3, RS=20/3) mice. (**H**) A ChIEF AAV was injected into the vSUB and optogenetically evoked EPSCs were recorded in the aBNST in male and female mice 24 hr after restraint stress. (**I, K**) Input-output representative traces (left), summary graph (middle), and slope (right) for EPSCs recorded in the aBNST of female (**I**; 0.262 pA, *P*>0.9999; 0.8 pA, *P*=0.9259; 1.29 pA, *P*=0.3001; 3.07 pA, \**P*=0.0262; 5.78 pA, \*\*\**P*=0.0004; 6.61 pA, \*\*\**P*=0.0006; slope, \**P*=0.0317; control n= 11/3, stress n=14/3) or male mice (**K**; 0.262 pA, *P=*0.9997; 0.8 pA, *P*=0.8669; 1.29 pA, *P*=0.2302; 3.07 pA, *P*=0.0662; 5.78 pA, \**P*=0.0240; 6.61 pA, \*\**P*=0.0025; slope, \**P*=0.0308; control n= 16/3, stress n=10/3). (**J, L**) Representative PPR traces (50ms) (left) and PPR measurements (right) from female (**J**; \**P*=0.0207, control n=13/3, stress n=13/3) or male (**L**; \*\*\**P*=0.0005, control n=16/3, stress n=14/3) mice. Data are represented as mean +/- SEM; means were calculated from the total number of cells. Numbers in the legend represent the numbers of cells/animals. Statistical significance was determined by a 2-way ANOVA or unpaired t-test.

### Stress exerts sex-specific effects on excitatory vSUB-aBNST synapses

Stress-sensitive vSUB BS cells overwhelmingly innervate aBNST cells, which raised the intriguing possibility that vSUB-aBNST synapses also exhibit functional adaptations to ARS. To test this, we injected AAV-ChIEF into the vSUB of P21 mice and electrophysiologically monitored light-evoked vSUB EPSCs from postsynaptic aBNST neurons. In females, stress weakened vSUB-aBNST excitatory synaptic strength, which was driven by decreased presynaptic release probability (Figures 5H-J). In contrast, stress strengthened vSUB-aBNST synapse in males which was driven by increased presynaptic release (Figures 5K-L). Together, in addition to the sex-specific changes to the vSUB microcircuitry, ARS causes sex-dependent changes in vSUB output to the aBNST; a projection that critically regulates anxiety-like behavior and corticosterone release.^14,16^

### Stress induces sex-specific changes to vSUB-aBNST pathway activity *in vivo*

To expand on our *ex vivo* findings, we employed fiber photometry to monitor *in vivo* calcium signals in aBNST-projecting vSUB cells, which is a correlative approach to link neuronal activity and behavior.^53^ To selectively monitor the calcium dynamics of aBNST-projecting vSUB neurons, of which are ∼80% BS (Figures 5D-E), we utilized an intersectional viral approach where we stereotaxically injected aBNST with a retrograde AAV2 encoding Cre recombinase and tdTomato (AAV_rg_-hSyn-Cre-P2A-tdTomato) and stereotaxically injected an AAV encoding a Cre-dependent calcium indicator (AAV_1_-hSyn-FLEx^loxP^-jGCaMP7f)^54^ into vSUB. We assessed activity in two ways. One, we monitored Ca^2+^ activity time-locked to movement initiation before (i.e., control conditions) and after stress and time-locked to struggle initiation during ARS (Figures 6A-E) to assess behavior-evoked Ca^2+^ activity. Second, we assessed neuronal activity during the recordings by measuring the number of events and the percentage of time spent above an activity threshold, as performed previously.^54–56^ In females, ARS increased the maximum Z-score amplitude, assessed during the window 10 seconds before and 20 seconds after a motion event, by ∼40% compared to control (Figure 6F), which represents increased neuron activity during struggling. After ARS, the Z-score amplitudes during movement were not different from the control period (Figure 6F). Although the Z-score amplitudes were significantly increased during movement in ARS, we did not observe a significant concomitant increase in the frequency of Ca^2+^ transients or in the percentage of time the activity of the defined cell population spent above threshold, which serves as a proxy for activity of the cells during the whole recording (Figures 6G-H).

**Figure 6.**
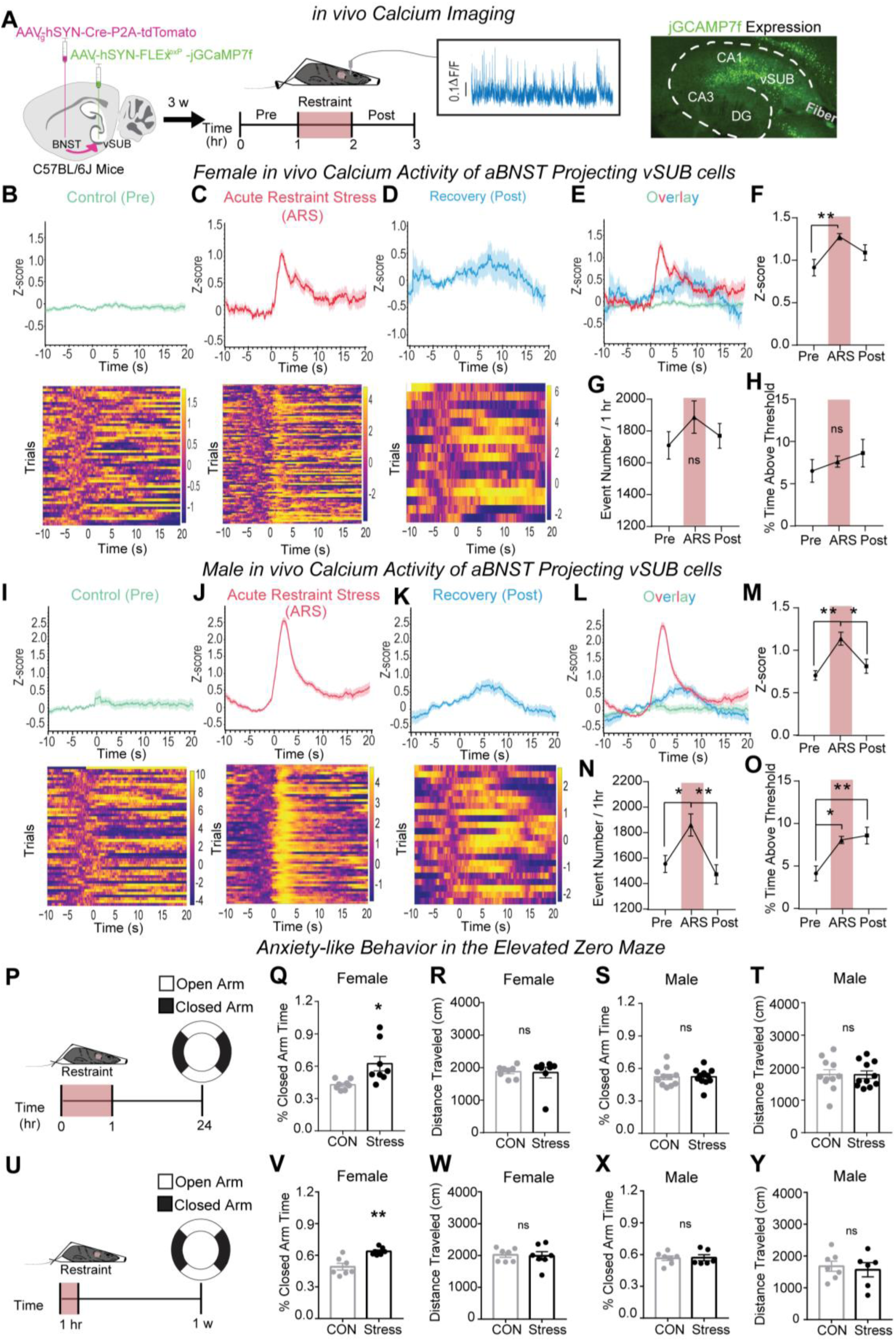
ARS induces sexually dimorphic responses in vSUB-aBNST *in vivo* calcium activities and anxiety-like behavior. (a) Schematic of dual virus injection and monitoring in vivo calcium activity of aBNST-projecting vSUB cells in control (pre), ARS, and stress recovery periods (post). (**B-D**, **I-K**) The representative traces and heatmaps of *in vivo* calcium activities time-locked to locomotor initiation during the control phase (**B** female; **I** male) and the stress recovery phase (**D** female; **K** male) or time-locked to struggle initiation (i.e. trunk movement) during the ARS procedure (**C** female; **J** male) in mice. (**E**,**L**) Overlay of representative traces for females (**E**; **B-D**) and males (**L**; **I-K**). (**F**,**M**)The amplitude of calcium events represented as the Z-score of calcium events in female (**f**; Pre vs ARS \*\**P*=0.0095; Pre vs Post P=0.2283; ARS vs Post P=0.1968; n=7) and male (**M**; Pre vs ARS \*\**P*=0.0029; Pre vs Post *P*=0.5494; ARS vs Post \**P*=0.0198; n=7) mice. (**G**,**N**) The average calcium event frequency during each phase in females (**G**; *P*=0.1210; n=7) and males (**N**; Pre vs ARS \**P*=0.0184; Pre vs Post *P*=0.6673; ARS vs Post \*\**P*=0.0039; n=7). (**H**,**O**) Overall population activity represented as the percent time above threshold in females (**H**; *P*=6835; n=7) and males (**O**; Pre vs ARS \**P*=0.0121; Pre vs Post \*\**P*=0.0053; ARS vs Post *P*=0.8916; n=7). (**P**) Male and female mice were restrained for 1 hr and 24 hrs later, anxiety-like behavior was assessed using the EZM. (**Q**,**S**) Anxiety-like behavior is conveyed as the percent time spent in the closed arm of the EZM for females (**Q**;\**P*=0.0233; control n=8, stress n=8) and males (**S**; *P*=0.9781; control n=11, stress n=11). (**R**,**T**) Distance traveled during the EZM test in females (**R**; *P*=0.1949) and males (**T**; *P*=0.7477). (**U**) Male and female mice were restrained for 1 hr and 1 w later, anxiety-like behavior was assessed using the EZM. (**V**,**X**) Anxiety-like for females (**V**;*P=0.0011; control n=7, stress n=7) and males (**X**; P=0.9081; control n=7, stress n=6). (**W**,**Y**) Distance traveled during the EZM test in females (**W**; P=0.8485) and males (**Y**; P=0.6807). Statistical significance was determined by a 1-way ANOVA or unpaired t-test. See also Figures S8.

In males, ARS increased the Z-score magnitude relative to the control period upon movement by ∼61%, which returned to baseline during the recovery period following ARS (Figure 6M). Distinct from females, we also observed a significant ∼20% increase in the frequency of Ca^2+^ transients (Figure 6N). Interestingly, these changes in Z-score and transient frequency were more significant in magnitude in males compared to females. Unlike females, the percent time above threshold of aBNST-projecting vSUB cells was significantly increased during ARS and persisted through the stress recovery period (Figure 6O). These findings are consistent with and provide insight into the impact of the ARS-induced shift in BS cell E/I balance 24-hrs post-stress (Figure 4D). Collectively, ARS increases the activity of aBNST-projecting vSUB cells when time-locked to struggle in both sexes. However, only activity during struggling is statistically significant in females while acute activity during struggling as well as the overall Ca^2+^ transient frequency and percent time above threshold are significant in males. Further, elevated cellular activity persists in males during the recovery period. Importantly, the changes we observed in the time spent above the activity threshold in males cannot be explained by gross changes in locomotion during the recordings (Figure S8).

### Stress causes anxiety-like behavior in females but not males

Given the remarkable sex differences in *ex vivo* synaptic transmission and in *in vivo* calcium imaging, we next examined whether ARS impacted behavior that models anxiety disorders, which has a sex-biased incidence. We used the elevated zero maze (EZM) to measure anxiety-like behavior 24-hrs post ARS and observed increased anxiety-like behavior in females but not males (Figure 6Q-T). Further, ARS did not alter locomotion in either sex. Consistent with the enduring synaptic adaptations in females (Figure S7D), anxiety-like behavior was still increased 1-week after ARS (Figures 6 U-Y). These data mirror clinical work that identify sex-specific vulnerability to stress disorders^1^ and preclinical work that indicate females are more likely to exhibit anxiety- or depressive-like behaviors after acute stressors.^57–59^ In parallel with the sex-specific changes in synaptic and cellular activity we uncovered after ARS, we demonstrate that ARS also produces a sex-dependent outcome in anxiety-like behavior that is in line with clinical and preclinical literature.

### Sex-specific mechanisms underlie stress-induced adaptations of PV-BS inhibition

To begin to understand the mechanisms that may contribute to these sex-specific synaptic and behavioral phenotypes after ARS, we examined the roles of systemic SNS and corticosterone signaling. In response to environmental stressors, the locus coeruleus (LC) releases norepinephrine (NE), which activates α and β adrenergic receptors (ARs), and the HPA axis signals the release of corticosterone, which activates glucocorticoid (GR) and mineralocorticoid (MR) receptors.^60^ Although understudied, vSUB cells robustly express ARs, MRs and GRs.^23–26^ To prevent SNS and corticosterone signaling, we pre-treated animals with propranolol (β-AR blocker) and phentolamine (α-AR blocker) or metyrapone (corticosterone synthesis inhibitor), respectively.

In female controls, pretreatment with AR blockers did not impact basal PV inhibition (Figures 7A-D). However, pretreatment with AR blockers 30 minutes before ARS prevented the significant ARS enhancement of PV-BS inhibition (Fig. 7E-H). Interestingly, inhibiting corticosterone synthesis before ARS did not prevent ARS augmentation of PV-BS inhibition (Figures 7I-L). Blocking sympathetic or corticosterone signaling did not alter the basal or post-ARS strength of PV-RS inhibition (Figure S9). Together, these data surprisingly indicate that SNS signaling plays a more critical role than corticosterone signaling in mediating the synapse-specific enhancement of PV-BS inhibition in female mice following ARS.

**Figure 7.**
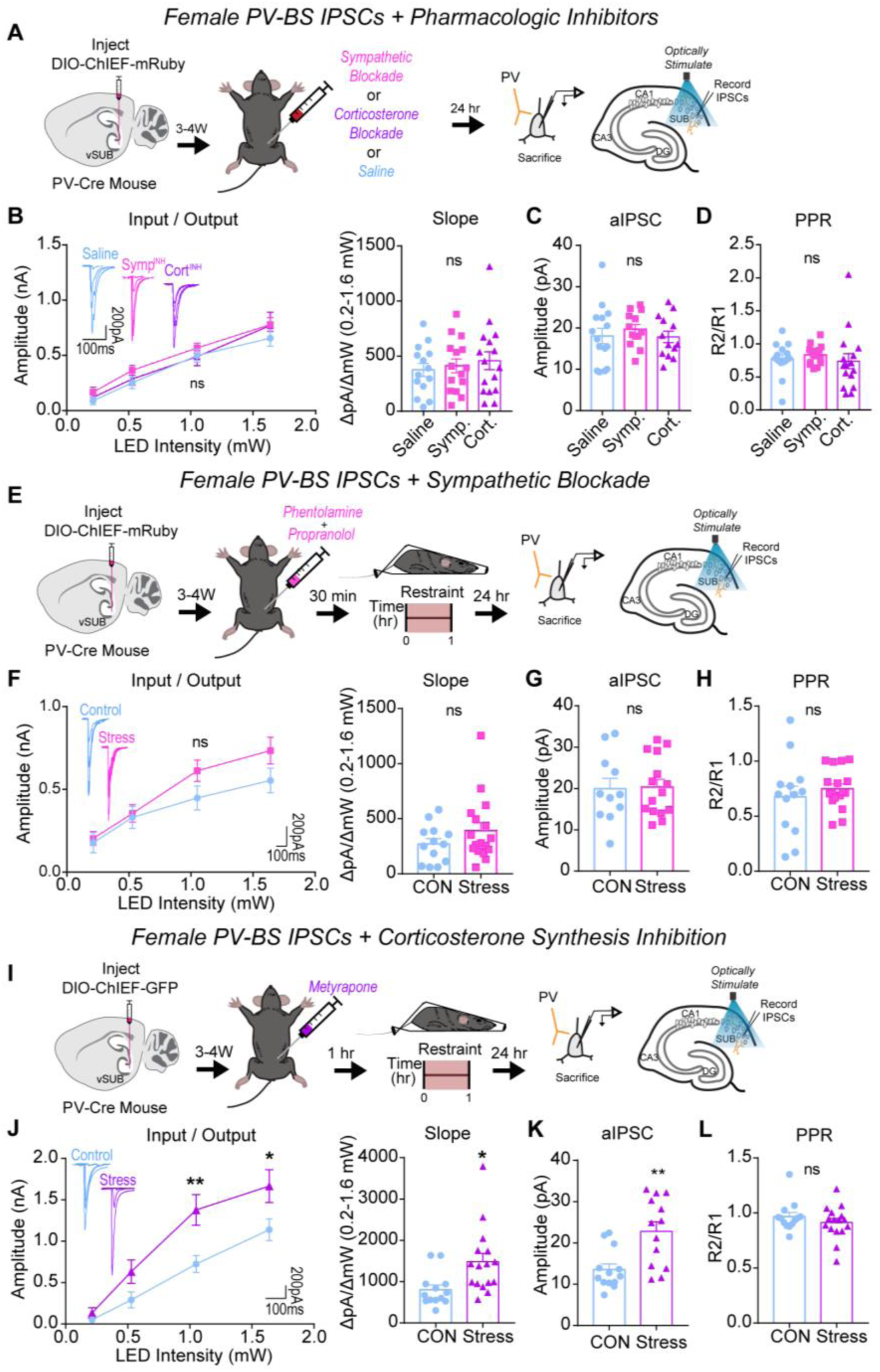
Systemic sympathetic signaling drives ARS-enhancement of PV-BS inhibition in females. (**A**) A Cre-dependent ChIEF AVV was injected into the vSUB of PV-Cre female mice and optogenetically evoked IPSCs from PV interneurons were recorded from BS cells 24-hr after saline, sympathetic block (symp), or corticosterone block (cort) pretreatment. (**B**) Input-output summary graph with representative traces (left) and slope (right) for IPSCs recorded in BS cells after drug controls (**B**; *P*=0.4011; slope, *P*=0.4583; saline n=14/3, symp n=15/3, cort n=16/3). (**C**) Strontium-mediated aIPSC amplitudes after optogenetic stimulation for BS cells (**c**; P= 0.6628; saline n=15/3, symp n=13/3, cort n=13/3). (**D**) PPR (50ms) measurements from BS cells (**D**; *P*=0.6986; saline n=14/3, symp n=15/3, cort n=16/3). (**E**,**I**) A Cre-dependent ChIEF AVV was injected into the vSUB of PV-Cre female mice, mice received a symp (**e**) or cort (**i**) pretreatment before ARS or underwent control conditions, and optogenetically evoked IPSCs from PV interneurons were recorded from BS cells 24-hr later. (f,j) Input-output summary graph with representative traces (left) and slope (right) for IPSCs in BS cells in symp (**E**; *P*=0.1973; slope, *P*=0.1960; control n=13/3, symp=17/4) or cort (**i**; 0.213 pA, *P*=0.9891; 0.518 pA, *P*=0.2942; 1.050 pA, \*\**P*=0.0044; 1.640 pA, \**P*=0.0320; slope, \**P*=0.0419 ; control n=13/3, cort n=16/3) studies. (**G**,**K**) Strontium-mediated aIPSC amplitudes from BS cells after symp (**G**; *P*=0.9042; control n=11/3, stress n=15/4) or cort (**K**; \*\**P*=0.0018, control n=13/3, cort n=13/3) studies. (**H**,**L**) PPR (50ms) measurements in BS cells in symp (**H**; *P*=0.4673, control n=13/3, symp n=17/3) or cort (**L**; *P*=0.3180, control n=13/3, cort n=16/3) studies. Numbers in the legend represent the numbers of cells/animals. Statistical significance was determined by a 1-way ANOVA, 2-way ANOVA, or unpaired t-test. See also Figures S9.

In male controls, pretreatment with AR or corticosterone blockers alone did not impact basal PV-BS inhibition (Figures 8A-D). Distinct from females, pretreatment with AR blockers did not prevent the ARS-mediated depression of inhibition at PV-BS synapses (Figures 8E-H). By contrast, and further highlighting the synaptic sex differences induced by ARS, the ARS-mediated depression of PV-BS synapses was absent when male mice were pretreated with metyrapone (Figure 8I-L). Similar to females, blocking sympathetic or corticosterone signaling before ARS did not alter PV-RS inhibition (Figure S10). In sum, our data implicate two distinct signaling pathways in the generation of ARS-induced adaptations in PV-BS inhibition. SNS signaling through ARs drives ARS-induced PV-BS inhibitory adaptations in females whereas corticosterone signaling plays a larger role in males.

**Figure 8.**
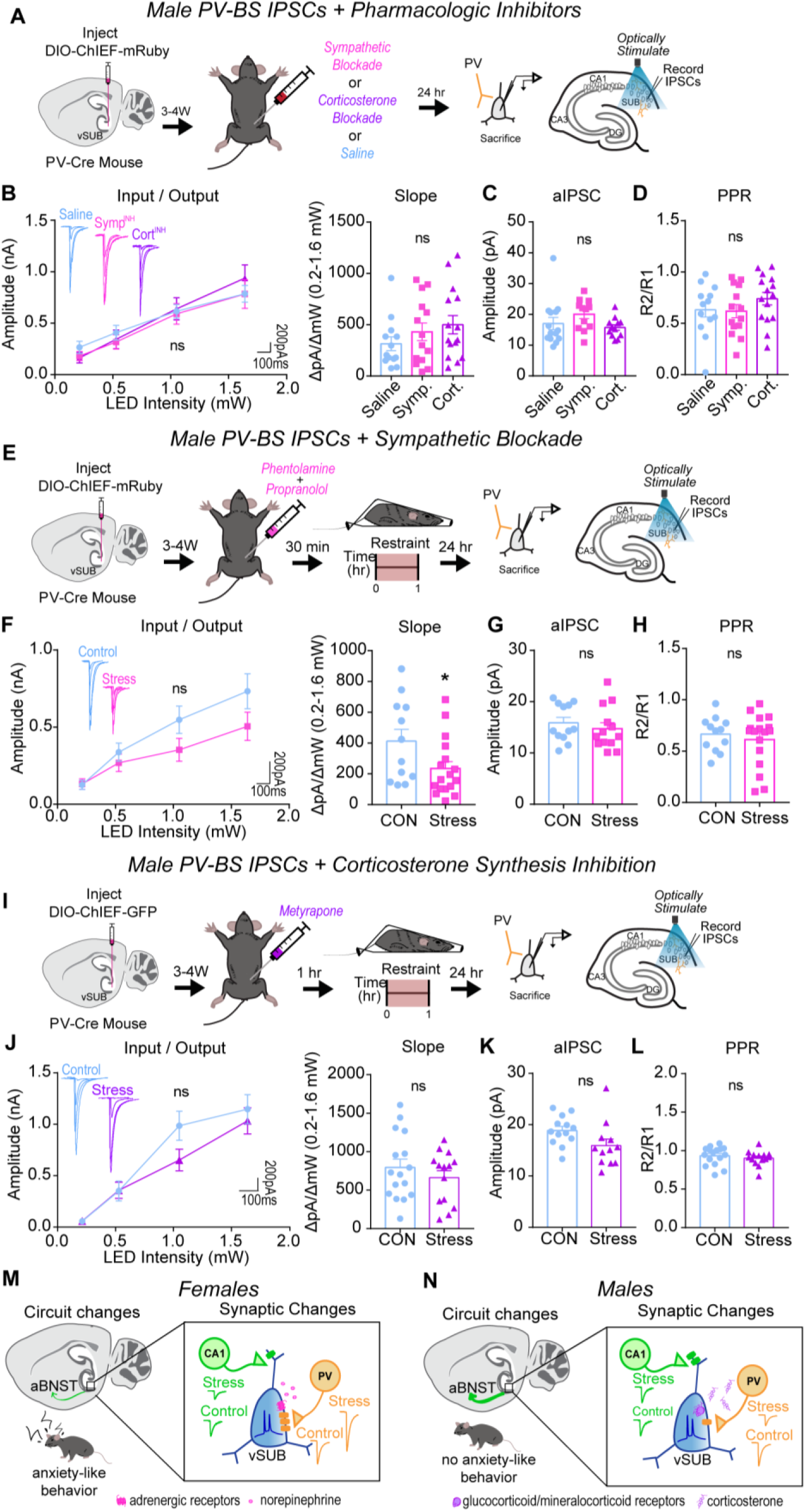
Systemic corticosterone signaling drives ARS-reduction of PV-BS inhibition in males. (**A**) A Cre-dependent ChIEF AVV was injected into the vSUB of PV-Cre male mice and optogenetically evoked IPSCs from PV interneurons were recorded from BS cells 24-hr after saline, sympathetic block (symp), or corticosterone block (cort) pretreatment. (**B**) Input-output summary graph with representative traces (left) and slope (right) for IPSCs recorded in BS cells after drug controls (**B**; *P*=0.8472; slope, *P*=0.2972; saline n=13/3, symp n=14/3, cort n=15/3). (**C**) Strontium-mediated aIPSC amplitudes after optogenetic stimulation for BS cells (**C**; P= 0.1645; saline n=14/3, symp n=12/3, cort n=12/3). (**D**) PPR (50ms) measurements from BS cells (**D**; *P*=0.3462; saline n=13/3, symp n=14/3, cort n=15/3). (**E**,**I**) A Cre-dependent ChIEF AVV was injected into the vSUB of PV-Cre female mice, mice received a symp (**E**) or cort (**I**) pretreatment before ARS or underwent control conditions, and optogenetically evoked IPSCs from PV interneurons were recorded from BS cells 24-hr later. (**F**,**J**) Input-output summary graph with representative traces (left) and slope (right) for IPSCs in BS cells in symp (**e**; *P*=0.1942; slope, \**P*=0.0448; control n=12/3, symp=17/3) or cort (**i**; *P*=0.3476; slope, *P*=0.3602 ; control n=16/3, cort n=14/3) studies. (**G**,**K**) Strontium-mediated aIPSC amplitudes from BS cells after symp (**G**; *P*=0.4761; control n=12/3, stress n=13/4) or cort (**K**; *P*=0.0563, control n=13/3, cort n=12/3) studies. (**H**,**L**) PPR (50ms) measurements in BS cells in symp (**H**; *P*=0.5388, control n=12/3, symp n=17/3) or cort (**L**; *P*=0.4401, control n=16/3, cort n=14/3) studies. (**M**,**N**) Summary graphs of synaptic, circuit, and behavioral findings after acute stress in females (**M**) and males (**N**). Data are represented as mean +/- SEM; means were calculated from the total number of cells. Numbers in the legend represent the numbers of cells/animals. Statistical significance was determined by a 2-way ANOVA or unpaired t-test. See also Figures S10.

## Discussion

Sex differences in brain circuitry represent potential drivers of sexually dimorphic behavioral responses to stress, with an emphasis on stress-related anxiety disorders.^57,61–63^ There is strong clinical and preclinical evidence that major acute stressors are particularly potent at inducing stress-related psychiatric disorders in females at greater rates than males,^6–8,57,64^ necessitating investigation into how each sex encodes acute stress. Here, we interrogated the circuitry of the vSUB, a major output of the vHipp that provides critical top-down regulation of the HPA axis.^16,65^ We recently demonstrated that vSUB exhibits basal sexual dimorphism,^17^ however, despite the known sex-specific responses to stress, sex as a biological variable has previously not been considered when studying the impact of stress on vSUB. Here, we provide a systematic synaptic interrogation of the impact of ARS on the vSUB local circuit and its output to aBNST with cell-type- and synapse-specific resolution in male and female mice.

ARS induced profound sex-specific synaptic adaptations to PV-mediated inhibition in vSUB. ARS potentiated inhibitory PV-BS synapses in females and depressed PV-BS inhibition in males. In females, these changes in PV-mediated inhibition were immediate and long-lasting, whereas in males, we only observed the depression of PV-BS synapses 24-hrs after ARS. Mechanistically, the potentiation of PV-BS synapses is primarily driven by SNS signaling through ARs in females. In males, depression of the same synapses is primarily caused by corticosterone signaling. Curiously, these sex-specific adaptations following ARS occurred on the same class of principal neurons and manifested via changes in postsynaptic strength which raises two key questions.

First, what is the consequence of these sex-specific stress adaptations observed locally within vSUB and in vSUB projections to aBNST? Our data indicate that vSUB output to the aBNST, a highly sexually dimorphic region integrated within stress and anxiety circuitry,^66–71^ is disproportionately dominated by BS cells. aBNST neurons are predominantly inhibitory and participate in feed-forward inhibition of downstream targets that govern the stress response and anxiety. In rodents exposed to aversive contexts, the activity of aBNST-projecting vSUB neurons decreases^15^ while experimentally increasing the activity of aBNST-projecting vSUB neurons decreases corticosterone during restraint stress and reduces anxiety-like behavior.^14,16^ Here, we found that ARS produces a diametrically opposite net effect on BS cell activity and in vSUB-aBNST output.

In vSUB BS cells, ARS shifts the E/I balance toward inhibition in females and excitation in males. At vSUB-aBNST synapses, ARS significantly decreases the strength of these synapses in females and increases synaptic strength in males (Figures 3-5). Together, the shift toward inhibition of BS cells and decreased strength of vSUB-aBNST synapses in females may contribute to a pathologic anxiety state. In contrast, the shift toward excitation of BS cells and potentiated vSUB-aBNST synapses in males may reflect a compensatory response to cope with an anxiogenic environment. Given the several distinct roles this projection contributes to through downstream signaling (i.e., corticosterone regulation, aversion, anxiety), it will be valuable to attain further functional granularity of postsynaptic aBNST neurons. There are at least five electrophysiologically distinct classes of BNST neurons and over 12 functionally distinct subregions^51^ and it will be important to characterize vSUB inputs onto these distinct classes of aBNST neurons to gain further resolution. Our work offers insight into sexually dimorphic input to aBNST and highlight that in addition to its inherent sex-specific organization, long-range inputs might also encode for the sexually dimorphic properties of aBNST.

Second, what are the molecular signaling pathways downstream of adrenergic and corticosterone signaling that might result in sex-specific synaptic adaptations following ARS? In females, the immediate onset and postsynaptic locus of the PV-BS synaptic adaptation are indicative of postsynaptic post-translational modifications and the sustained adaptation is indicative of changes in gene expression. Adrenergic signaling, largely in response to NE released from the LC, mediates the 24-hr effects of ARS (Figure 7) and can drive rapid changes in synaptic strength by triggering the phosphorylation of GABA receptors^72–74^ and gephyrin^75^ and can produce long-lasting changes by altering gene expression.^76^ Future studies are needed to determine how corticosterone signaling enables the delayed and transient plasticity of PV-BS synapses in males (Figure 8). Our sex-specific involvement of the SNS and corticosterone signaling agree with clinical work indicating that autonomic dysfunction is more common in females with generalized anxiety disorder^77,78^ and males with major depression disorders more frequently exhibit cortisol hyperactivity.^79^ Further, preclinical work has identified pronounced sexual dimorphism in LC connectivity, NE activity during puberty, and corticotropin-releasing-factor signaling that may contribute to sex-specific outcomes in stress-related psychiatric disease pathology, highlighting these as appealing signaling pathways for further examination.^80^

In addition to the sex-specific adaptations of PV-BS synapses and vSUB-aBNST output, we found that basal excitatory transmission and LTP at vCA1-BS synapses were impaired following ARS in females but unaltered in males (Figures 1, 2, S3, S4). Basal transmission at vCA1-BS synapses in females was reduced due to decreased postsynaptic strength, however, LTP at vCA1-BS synapses manifests presynaptically due to enhanced release probability. α2 and β2 ARs and MRs are localized in the pre- and post-synaptic membrane and activation of these presynaptic receptors facilitate presynaptic release, making them unlikely candidates as drivers of the impaired vCA1-BS LTP we see here. However, it is important to note that studies of presynaptic ARs and MRs did not include sex as a biological variable.^25,81,82^ It will be important to identify the signaling pathway involved in preventing presynaptic LTP in female vSUB. Our findings contrast with studies using different models of acute or chronic stress in males that alter LTP,^25,83,84^ thus different physiological stressors appear to be encoded uniquely at hippocampal synapses and it will be important to continue studying how varying stressors impact brain circuitry.

We found that ARS induced lasting anxiety-like behaviors in female but not male mice. Although anxiety-like behaviors have been reported in male rodents following acute stress, the stress paradigm used differed.^59,85,86^ Importantly, the anxiety-like behavior in females is consistent with our functional analysis in vSUB and aBNST. Curiously, when we monitored network activity *in vivo*, female vSUB-aBNST calcium transient amplitudes responded to ARS but this response diminished during the recovery period. Males had robust activation of vSUB-aBNST cells that persisted through the post-stress recovery period. Additionally, the onset of neural activity in both sexes coincided with the onset of struggling. Struggling activity is interpreted as active coping and has been attributed to a neural circuit comprised of the insular cortex and BNST.^87^ vSUB may participate in this coping mechanism and may be downstream of insular cortex and upstream of BNST or represent a parallel pathway to promote the struggle behavior. This would suggest that perhaps vSUB is situated to provide top-down regulation of the HPA axis and participate in stress-coping mechanisms. Our data provide insight into the behavioral impact of our functional analyses and may elaborate on circuits responsible for active coping during stress.

Despite the clinical impetus to consider sex as a biological variable when studying models of stress, nearly all preclinical work has been conducted solely on male rodents.^88^ Here, we focused on sex differences in brain circuitry rather than circulating hormones. Both sexes have circulating levels of estrogen and testosterone, with estrogen fluctuations over a 4-day estrous cycle in female rodents and diurnal testosterone fluctuations with incredible individual variability of up to 20-fold differences between males.^89–91^ Although we did not track estrous cycle or testosterone levels here, we observed higher variability in males on basal excitatory, basal inhibitory, and activity-dependent excitatory synaptic strengths. The high individual variability in males is consistent with others^92,93^ and may be explained by testosterone levels and/or social hierarchy as the mice were group-housed,^94^ but more work is necessary to determine this. Our findings here provide strong evidence to evaluate sex as a biological variable when modeling diseases with sexually dimorphic features and illustrate striking sex differences in how acute stress is internalized in a cell-type, synapse-, and behavior-specific manner.

## Author contributions

C.N.M performed AAV injections, electrophysiology experiments, behavior experiments, and performed data analyses. Y.L. performed the fiber photometry data acquisition and analysis. K.T.B. performed fiber photometry analysis and aided in fiber photometry data interpretation. C.N.M. and J.A. are responsible for the study conception, experimental design, data interpretation, and manuscript generation. All authors provided input on the manuscript.

## Acknowledgements

This work was supported by the National Institutes of Health Grants R01MH116901 (to J.A.), F30DA057053 (to C.N.M.), 5T32GM007635 (to C.N.M.), and 5T32GM008497 (to C.N.M.). We thank Hannah Polatsek for her contributions to mouse behavioral data collection. We also thank members of the Aoto lab for thoughtful discussions.

## Methods

### Lead contact

For further information and requests for resources, data, and reagents should be directed to and will be fulfilled by the lead contact, Jason Aoto.

### Material availability

Plasmids generated in this study are available upon request.

### Data and code availability

- All original code and fiber photometry data have been deposited at Github or Zenodo, respectively, and is publicly available as of the date of publication. Accession codes are listed in the key resources table.
- Any additional information required to reanalyze the data reported in this paper is available from the lead contact upon request.

### Experimental model and subject details

For all electrophysiology experiments, male and female mice were bred and group-housed (2-5) in ventilated cages enriched with cotton nestlets on a 12-h light/dark cycle at the University of Colorado Anschutz. Transgenic PV-IRES-Cre mice (Jackson Laboratories, Stock No: 008069) breeders were kindly provided by Dr. Diego Restrepo and Gt(ROSA)26Sortm9(CAG-tdTomato)Hze mice were purchased from the Jackon Laboratory (“Ai9”: Jax 008909). All experimental mice were maintained on congenic B6;129 or B6.Cg mixed genetic backgrounds. Mice were given food and water *ad libitum* and maintained at 35% humidity, 21-23C, on a 12/12 light/dark cycle in a dedicated vivarium. Mice were genotyped in-house, and the sex of the animal was determined by external genitalia. All procedures were conducted in accordance with guidelines approved by Administrative Panel on Laboratory Animal Care at University of Colorado, Anschutz School of Medicine, accredited by Association for Assessment and Accreditation of Laboratory Animal Care International (AAALAC) (00235).

All fiber photometry experiments took place at the University of California, Irvine, and were in accordance with the National Institutes of Health Guidelines for Animal Care and Use. 8-week-old male and female C57BL/6J mice were purchased from the Jackson Laboratory (“B6”: Jax 000664) and housed in separate cages according to sex. Mice were housed in standard ventilated cages with corn cob bedding and two cotton cots for environmental enrichment. Lights were on a 12-hour on/12-hour off cycle (7:30 am-7:30 pm), and room temperature (22°C +/- 2°C) and humidity (55-65%) were controlled. Mice had free access to food and water. Mice were housed at the Gillespie Neuroscience Research Facility. For all studies, the experimental group assignment (control vs stress) was counterbalanced across littermates and age-matched. All experiments were conducted in males and females and the data were stratified by sex.

## Method details

### Restraint stress procedure

Adult animals were habituated to handling for 3-5 days (5min/day) prior to stress or control procedures. Handling mice consisted of mice freely moving between the experimenters’ hands and mice were scooped rather than picked up from their tails to avoid undesired stress. Acute restraint stress was administered for 1hr using commercially available plastic cones (Braintree Scientific Inc) fastened with a twisty tie in a fresh cage. Mice were completely unable to move all limbs during the restraint procedure. Control mice moved freely for 1-hr in a fresh cage. Control or stress procedures all occurred between 0800-1000. Control and stress animals were returned to their home cage or sacrificed for electrophysiology experiments immediately after either procedure. For pharmacologic intervention studies, adrenergic blockage was achieved with propranolol (10 mg/kg in saline; i.p.; Sigma) and phentolamine (15 mg/kg in saline; i.p.; Sigma) 30 min prior to restraint or control procedures. Corticosterone synthesis was inhibited by administering metryapone (75-100 mg/kg in DMSO, i.p; Cayman Chemical) 1-hr before restraint or in untreated animals. Doses for adrenergic blockade^95,96^ and corticosterone synthesis inhibitors^97,98^ were determined by previous literature. All injectable solutions were prepared fresh on the morning of.

### Elevated zero maze

24-hrs or 1 week after stress or control conditions, adult mice (P42-60) were tested in the elevated zero maze (EZM) during their light phase of the light/dark cycle to assess anxiety-like behavior.^99^ Mice were accustomed to the behavior room for at least 1-hr before testing. The mouse test order was counterbalanced by pretreatment and sex. White noise was played during the habituation and experimentation period to minimize unwanted disruption. Experimenters remained outside of the room during the 10 min of behavior recording. The EZM consisted of 4 quadrants with a 40 cm inner diameter, 2 closed-arm sections, and 2 open-arm sections as illustrated in Fig 6. Mouse movement was tracked with Ethovision software. The percentage of time in each arm and total distance traveled were analyzed from the first 5 min of the EZM because this period represents the most robust anxiogenic response period^100^.

### Stereotactic viral injections

Stereotactic injections were performed on P21-22 mice (8-14g) for electrophysiological experiments. Animals were induced with 4% isoflurane, maintained at 1-2% isoflurane, and then head fixed to a stereotactic frame (KOPF). Small holes were drilled into the skull and 0.1-0.2 μL solutions of adeno-associated viruses (AAVs) were injected with glass micropipettes into the brain at a rate of 10 μL/hr using a syringe pump (World Precision Instruments), consistent with standard protocols^17^. Coordinates (in mm) for ventral subiculum (vSUB) were (A-P/M-L/D-V from Bregma and relative to pia): -3.41, +/- 3.17, -3.45. Coordinates (in mm) for anterior ventral bed nucleus of the stria terminalis (avBNST) were: +0.15, +/-1.1, -4.5. All AAVs used in this study were packaged in-house: AAV DJ-hSYN-DIO_loxp_-ChIEF-mRuby, AAV DJ-hSYN-ChIEF-mRuby, and AAV2-Retro-mRuby, consistent with standard protocols^17,36^. All AAV vectors were created from an empty cloning vector with the expression cassette is as follows: left-ITR of AAV2, human synapsin promotor, 5’ LoxP site, multiple cloning site, 3’ LoxP site, and right ITR. All AAV plasmids were co-transfected with pHelper into HEK293T cells. Cells were then harvested and lysed 72-hrs post transfection, and the virus was harvested from the 40% iodixanol fraction. Finally, the virus was concentrated in 100K MWCO ultracon filter.

### *Ex vivo* whole-cell electrophysiology

Adult (P42-60) mice were deeply anesthetized with isoflurane, decapitated, and their brains were rapidly dissected, consistent with standard protocols.^17,36^ Brains were removed and 300 μm horizontal slices containing the vSUB or 250 μm coronal slices containing the avBNST were sectioned with a vibratome (Leica VT1200) in ice-cold high-sucrose cutting solution containing (in mM) 85 NaCl, 75 sucrose, 25 D-glucose, 24 NaHCO_3_, 4 MgCl_2_, 2.5 KCl, 1.3 NaH_2_PO_4_, and 0.5 CaCl_2_. Slices were transferred to 31.5°C oxygenated artificial cerebral spinal fluid (ACSF) containing (in mM) 126 NaCl, 26.2 NaHCO_3_, 11 D-Glucose, 2.5 KCl, 2.5 CaCl_2_, 1.3 MgSO_4_-7H_2_O, and 1 NaH_2_PO_4_ for 30 min, then recovered at room temperature for >1hr before recordings. Recordings were made at 29.5°C ACSF, with 3-5 MΩ patch pipettes, and cells were voltage-clamped at -70mV. Pyramidal neurons in the vSUB were visualized with a BX51 microscope with a 40x dipping objective collected on a Hamamatsu ORCA-Flash 4.0 V3 digital camera with an IR bandpass filter. The identity of pyramidal neurons (regular (RS) versus burst spiking (BS)) was determined by current-clamping and injecting a 500ms depolarizing current in 50 pA steps, as previously described^17^. Cells that exhibited burst firing upon suprathreshold current injection (2-4 spikes with ∼10ms inter-spike-interval) were considered BS and those that did not were labeled RS.

To measure inhibitory currents, NBQX (10 μM) and D-AP5 (50 μM) were added to the ACSF, and a high-chloride internal solution containing (in mM) 95 K-gluconate, 50 KCl, 10 HEPES, 10 Phosphocreatine, 4 Mg_2_-ATP, 0.5 Na_2_-GTP, and 0.2 EGTA was used. To measure excitatory currents, picrotoxin (100 μM) was added to the ACSF, and a low-chloride internal solution containing (in mM) 137 K-gluconate, 5 KCl, 10 HEPES, 10 Phosphocreatine, 4 Mg_2_-ATP, 0.5 Na_2_-GTP, and 0.2 EGTA was used. For strontium experiments, CaCl_2_ was replaced with 2.5 mM SrCl_2_ in the recording ACSF. To assess intrinsic excitability of neurons, cells were current-clamped and given −100 to +400 pA current injections in 20-50 pA steps. To optogenetically stimulate ChIEF-expressing fibers in the vSUB or BNST, slices were illuminated with 470 nm LED light (ThorLabs M470L2-C1) for 3 ms through the 40x dipping objective located directly over the recorded cell. With an illumination area of 33.18mm^2^ the tissue was excited with an irradiance of 0.006 to 0.17 mW/mm^2^. To electrically stimulate pyramidal cells, a bipolar nichrome stimulating electrode was made in-house and placed in the alveus/stratum oriens border of vCA1 and controlled by a Model 2100 Isolated Pulse Stimulator (A-M System, Inc.)^36^. To induce long-term potentiation of vCA1-vSUB synapses, we used a standard protocol of four 100 Hz tetani separated by 10 s.^21,36^

### Analysis of electrophysiology recordings

All recordings were acquired using Molecular Devices Multiclamp 700B amplifier and Digidata 1440 digitizer with Axon pClamp™ 9.0 Clampex software, lowpass filtered at 2 kHz and digitized at 10-20 kHz. Evoked PSC peak amplitudes from each recording were identified using Axon™ pClamp10 Clampfit software. To collect input/output curves, 8 sweeps (0.1 Hz) were averaged at each stimulation intensity. All final postsynaptic currents (PSC) values are displayed as the absolute value. The input/output slope was calculated using the SLOPE function in Microsoft Excel: (amplitude range/intensity range) from the linear range of the curves. The release probability was obtained over 10 sweeps (0.1 Hz) at each inter-stimulus interval (ISI) and was assessed by measurements of paired-pulse ratios (PPRs) at 25-1000 ms ISI. The PPR was measured by dividing the average PSC amplitude evoked by the second stimulus by the average PSC amplitude evoked by the first stimulus (R2/R1). For strontium experiments, the slices were stimulated over 15 sweeps (0.25 Hz) and the amplitudes of asynchronous PSC events that occurred within a 300 ms window immediately following the phasic PSC were analyzed using Clampfit event detection software.^17^

### Fiber photometry procedure

8-week old adult mice were injected with 200 μL of AAVrg-hSyn-Cre-P2A-tdTomato (2.3×10¹³ vg/mL) into the aBNST at the coordinates AP +0.1; ML 0.8; DV -4.5 (in millimeters relative to the Bregma, midline, or dorsal surface of the brain) at a rate of 1.6 nL/sec. During the same surgery, 500 μL of AAV1-hSyn-FLEx^loxP^-jGCaMP7f (1.2×10¹³ vg/mL) was injected into the vSub at coordinates AP -4.04; ML 3.4; DV -3.5. The glass capillary was left in place for 5 minutes following injection before removal. After completing the two injections, a 400 μm diameter, 0.39NA, 3mm-long optical fiber was implanted directly over the vSub. The implantation site was targeted at 0.1mm dorsal to the injection site. Implants were secured to the skull with metal screws (Antrin Miniature Specialists), Metabond (Parkell), and Geristore dental epoxy (DenMat). Mice recovered for at least 2 weeks before experiments began.

Mice were handled for three days before the start of recordings to reduce the novelty of handling. On recording days, recordings were performed from 8:30 a.m. to 12:00 p.m. to match potential circadian effects in the electrophysiology experiments. Prior to performing recordings, the top of the ferrule was cleaned using a 70% alcohol solution. Following this, each of the three phases was recorded, sequentially, each with a recording time of one hour. At the end of each one-hour recording session, the mice were moved to a new cage to avoid the influence of odor from the previous session. Throughout the entire three-hour recording period, the mice had no access to water or food.

During the pre-restraint stage, following patch cable connection, mice were placed into a clean cage and allowed to move freely for one hour. Mice were disconnected from the patch cable after this recording was complete. During the restraint stage, mice were enclosed in a conical plastic bag with open holes on the right top of their heads. The holes were only large enough to allow the ferrule to pass through. After enclosing the mouse, we wrapped the excess plastic portion around the mouse’s body and secured the wrap with tape and staples, ensuring that the mouse was able to comfortably breathe. After laying the mouse flat into the new cage, the mouse was again connected to the patch cord, and recording was performed for one hour. Mice were again disconnected from the patch cable after this recording was complete. During the post-restraint recovery phase, mice were freed from the restraints, at which time the mice were again connected to the patch cord, and another hour of recording was performed, as in the pre-restraint stage.

### Analysis of fiber photometry recordings

Fiber photometry recordings were made using previously described equipment.^101^ In brief, 465 nm and 405 nm excitation light were controlled via a RZ5P real-time processor (Tucker Davis Technologies) using Synapse software and were used to stimulate Ca^2+^-dependent and isosbestic emission, respectively. All optical signals were bandpass filtered with a Fluorescence Mini Cube FMC4 (Doric) and were measured with a femtowatt photoreceiver (Newport). Signal processing was performed with MATLAB (MathWorks Inc.). For the definition of timestamps for motion, during pre- and post-restraint sessions the motion of the mouse was tracked using biobserve software. Following the video recording, a custom Python script calculated velocity of the mouse by taking the distance the center body coordinate of the mouse moved from frame to frame. The velocity threshold for a motion event was a sustained event with velocity higher than 12.5 cm/s. For the restraint phase, as mice could not locomote within their restraint, motion was defined visually by the observer as events where the mouse clearly moved their trunks as well as their heads. Events where the mouse only shook their heads were not counted.

All traces from individual mice were manually screened to exclude individuals with traces that contained high levels of noise and/or unclear signals. Signals were first motion corrected by subtracting the least squares best fit of the control trace to the calcium signal. Data points containing large motion artifacts were then manually removed. To assess neural activity, we quantified the percent of time spent above the threshold, which was set at 2.91 times the median absolute deviation (MAD) of each day’s recording, a value that equates to the 95% confidence interval for Gaussian data.^102^ We then used the Guppy script to compute the amplitude and frequency for each mouse and trace.^103^

### Quantification and statistical analyses

All statistical analyses were performed in Prism 7 (GraphPad). For two-group comparisons, we used a student-paired or unpaired 2-tailed t-test where appropriate. To estimate the relationship between locomotor activity and calcium signal in fiber photometry experiments, a simple linear regression was used. For multiple group comparisons, we used a one-way ANOVA or two-way ANOVA where appropriate. Repeated-measures ANOVA was also used when appropriate. Šídák *post-hoc* comparisons were performed following ANOVAs. All experiments were replicated in at least 3 animals/condition/sex and littermate matched across groups. Whenever possible, the experimenter was blinded to animal sex, treatment, and cell identity during analysis. Cells of low quality were excluded from the electrophysiology analysis (i.e, unstable baseline, access resistance >10% of the membrane resistance). Statistical tests used for each experiment and exact p values are present in the figure legends. In figures, *p<0.05, **p<0.01, ***p<0.001, and ****p<0.0001, ns p>0.05. Samples can be found within figures or in figure legends and are notated: n/N= cell number/animal number. Averages are graphically expressed as arithmetic mean +/- SEM.

## Supplemental figure legends

**Supplemental Figure 1 (Relates to Figure 1).** Stress ablates CA1-vSUB presynaptic LTP in BS cells of females. (**A**,**E**) Summary graph of LTP experiments in BS (**A**) or RS (**E**) cells in females. (**B**,**F**) Representative LTP traces in BS (**B**) or RS (**F**) cells. (**C**,**G**) LTP magnitude from the last 10 minutes of LTP recording in BS (**C**; \**P*=0.044; control n=11/6, stress n=10/4) or RS (**G**; *P*=0.5790; control n=10/6, stress n=13/5) cells.(**D**,**H**) Pre and post LTP PPR measurements in BS (**D**; control \*\*\**P*=0.008; stress *P*=0.2826; control n=8/6, stress n=7/4) or RS (**H**; control *P*=0.1970; stress *P*=0.2602; control n=9/6, stress n=9/5) cells. Data are represented as mean +/- SEM; means were calculated from the total number of cells. Numbers in the legend represent the numbers of cells/animals. Statistical significance was determined by a 1-way ANOVA, 2-way ANOVA, unpaired t-test, or paired t-test.

**Supplemental Figure 2 (Relates to Figure 2).** Stress does not alter CA1-vSUB LTP in males. (**A**,**E**) Summary graph of LTP experiments in BS (**A**) or RS (**E**) cells in males. (**B**,**F**) Representative LTP traces in BS (**B**) or RS (**F**) cells. (**C**,**G**) LTP magnitude from the last 10 minutes of LTP recording in BS (**C**; *P*=0.5539; control n=11/6, stress n=10/6) or RS (**G**; *P*=0.8880; control n=10/7, stress n=11/7) cells.(**D**,**H**) Pre and post LTP PPR measurements in BS (**D**; control \**P*=0.0353; stress \**P*=0.0151; control n=11/6, stress n=10/6) or RS (**H**; control *P*=0.3146; stress *P*=0.1656; control n=7/6, stress n=8/6) cells. Data are represented as mean +/- SEM; means were calculated from the total number of cells. Numbers in the legend represent the numbers of cells/animals. Statistical significance was determined by a 1-way ANOVA, 2-way ANOVA, unpaired t-test, or paired t-test.

**Supplemental Figure 3 (Relates to Figure 3).** Stress does not impact the intrinsic excitability of vSUB cell types of females. (**A**) (Left) Action potential firing frequency plotted by current injected with representative responses in BS cells to +260pA and (right) the minimum amount of current required to fire BS cells (Intrinsic excitability *P*=0.2884; rheobase, *P*=0.5594; control n=29, stress n=29). (**B**) (Left) Action potential firing frequency plotted by current injected with representative responses in RS cells to +260pA and (right) the minimum amount of current required to fire RS cells (intrinsic excitability *P*=0.3778; rheobase *P*=0.5594; control n=15, stress n=16). (**C**) (Left) Action potential firing frequency plotted by current injected with representative responses in tdTomato+ PV cells to +260pA and (right) the minimum amount of current required to fire PV cells in PVAi9 mice (intrinsic excitability *P*=0.1972; rheobase, *P*=0.8159; control n=14, stress n=20). Data are represented as mean +/- SEM; means were calculated from the total number of cells. Numbers in the legend represent the numbers of cells/animals. Statistical significance was determined by a 2-way ANOVA or unpaired t-test.

**Supplemental Figure 4 (Relates to Figure 4).** Stress reduces the intrinsic excitability of PV interneurons in the vSUB of males. (**A**) (Left) Action potential firing frequency plotted by current injected with representative responses in BS cells to +260pA and (right) the minimum amount of current required to fire BS cells (Intrinsic excitability *P*=0.1140; rheobase *P*=0.1739; control n=19/4, stress n=27/4). (**B**) (Left) Action potential firing frequency plotted by current injected with representative responses in RS cells to +260pA and (right) the minimum amount of current required to fire RS cells (Intrinsic excitability *P*=0.5848; rheobase *P*=0.5772; control n=18/3, stress n=20/3). (**C**) (Left) Action potential firing frequency plotted by current injected with representative responses in tdTomato+ PV cells to +260pA and (right) the minimum amount of current required to fire PV cells in PVAi9 mice (0-120pA *P*>0.9999; 140pA *P*=0.997; 160pA *P*=0.8984; 180pA *P*=0.3150; 200pA \*\**P*=0.0030; 220pA \*\**P*=0.0017; 240pA \**P*=0.0101; 260 pA \**P*=0.0384; 280pA \**P*=0.0384; 300pA *P*=0.0901; 320pA *P*=0.3583; 340pA *P*=0.8034; 360pA *P*=0.9319; 380pA *P*=0.9602; 400pA *P*=0.9996; rheobase \**P*=0.0144; control n=13/3, stress n=14/3). Data are represented as mean +/- SEM; means were calculated from the total number of cells. Numbers in the legend represent the numbers of cells/animals. Statistical significance was determined by a 2-way ANOVA or unpaired t-test.

**Supplemental Figure 5 (Relates to Figure 3).** Stress immediately enhances PV-vSUB-BS inhibition in females. (**A**) A Cre-dependent ChIEF AAV was injected into vSUB of PV-Cre female mice and optogenetically evoked IPSCs from PV interneurons were recorded from BS or RS cells immediately after 1-hr of ARS. (**B**,**E**) Input-output representative traces (left), summary graph (middle), and slope (right) for IPSCs recorded in BS (**B**; 0.213 pA, *P*=0.2616; 0.518 pA, \**P*=0.0276; 1.050 pA, \*\**P*=0.0015; 1.640 pA, \*\*\*\**P*<0.0001; **slope, *P*=0.0011; control n=14/3, stress n=16/3) or RS cells (**E**; 0.213 pA, *P*=0.8249; 0.518 pA, *P*=0.5096; 1.050 pA, *P*=0.2844; 1.640 pA, *P*=0.1869; slope, *P*=0.2217; control n=13/3, stress n=12/3). (**C**,**F**) Representative traces of strontium-mediated aIPSCs after optogenetic stimulation (left) and aIPSC amplitude (right) for BS (**C**; \*\**P*=0.0017; control n=16/3, stress n=14/3) or RS cells (**F**; *P*=0.1053; control n=11/3, stress n=9/3). (**D**,**G**) Representative PPR traces (50ms) (left) and PPR measurements (right) from BS (**D**; *P*=0.2770; control n= 14/3, stress n= 16/3) or RS cells (**G;** *P*=0.6621; control 14/3, stress n=12/3). Data are represented as mean +/- SEM; means were calculated from the total number of cells. Numbers in the legend represent the numbers of cells/animals. Statistical significance was determined by a 2-way ANOVA or unpaired t-test.

**Supplemental Figure 6 (Relates to Figure 4).** Stress does not have immediate effects on PV-vSUB inhibition in males. (**A**) A Cre-dependent ChIEF AAV was injected into vSUB of PV-Cre male mice and optogenetically evoked IPSCs from PV interneurons were recorded from BS or RS cells immediately after 1-hr of ARS. (**B**,**E**) Input-output representative traces (left), summary graph (middle), and slope (right) for IPSCs recorded in BS (**B**; *P*=0.1605; slope, *P*=0.0011; control n=14/3, stress n=18/3) or RS cells (**E**; *P*=0.9132; slope, *P*=0.1130; control n=12/3, stress n=11/3). (**C**,**F**) Representative traces of strontium-mediated aIPSCs after optogenetic stimulation (left) and aIPSC amplitude (right) for BS (**C**; \*\**P*=0.0017; control n=13/3, stress n=13/3) or RS cells (**F**; *P*=0.4484; control n=10/3, stress n=9/3). (**D**,**G**) Representative PPR traces (50ms) (left) and PPR measurements (right) from BS (**D**; *P*=0.3285; control n=14/3, stress n=18/3) or RS cells (**G;** *P*=0.2980; control 12/3, stress n=11/3). Data are represented as mean +/- SEM; means were calculated from the total number of cells. Numbers in the legend represent the numbers of cells/animals. Statistical significance was determined by a 2-way ANOVA or unpaired t-test.

**Supplemental Figure 7 (Relates to Figures 3-4).** The impact of stress on PV-vSUB inhibition is lasting in females, but transient in males. (**A**) A Cre-dependent ChIEF AAV was injected into vSUB of PV-Cre mice and optogenetically evoked IPSCs from PV interneurons were recorded from BS or RS cells 1-week after 1-hr of ARS. (**B**) Representative firing patterns from BS (blue) and RS (red) cells in response to 150pA current. (**C**,**G**) Proportion of cells patched that are RS or BS in females (**C**; stress BS n=31/3, RS n=1/3; control BS n=26/3, RS n=22/3) or males (**G**; stress BS n=28/3, RS n=25/3; control BS n=27/3, control n=25/3). (**D**) Input-output representative traces (left), summary graph (middle), and slope (right) for IPSCs recorded in female BS cells (**B**; 0.213 pA, *P*=0.9812; 0.518 pA, *P*=0.4146; 1.050 pA, \**P*=0.0153; 1.640 pA, \*\*\**P=*0.0002; slope, \**P*=0.0196; control n=15/3, stress n=16/3). (**E**) Representative traces of strontium-mediated aIPSCs after optogenetic stimulation (left) and aIPSC amplitude (right) for female BS (\*\**P*=0.0088; control n=12/3, stress n=12/3) cells.(**D**,**G**) Representative PPR traces (50ms) (left) and PPR measurements (right) from BS (**D**; *P*=0.4702; control n= 15/3, stress n= 15/3). (**H**,**K**) Input-output representative traces (left), summary graph (middle), and slope (right) for IPSCs recorded in male BS (**H**; *P*=0.8751; slope *P*=0.6435; control n=14/3, stress n=15/3) or RS (**K**; *P*=0.8714; slope *P*=0.9537; control n=15/3, stress n=13/3) cells. (**i**,**l**) Representative traces of strontium-mediated aIPSCs after optogenetic stimulation (left) and aIPSC amplitude (right) for male BS (**I**; *P*=0.4757; control n=13/3, stress n=13/3) or RS (**l**; *P*=0.6625; control n=13/3, stress n=13/3) cells.(**J**,**M**) Representative PPR traces (50ms) (left) and PPR measurements (right) from BS (**J**; *P*=0.2635; control n= 16/3, stress n= 15/3) or RS (**M**; *P*=0.0955; control n=13/3, stress n=13/3) cells. Data are represented as mean +/- SEM; means were calculated from the total number of cells. Numbers in the legend represent the numbers of cells/animals. Statistical significance was determined by a 2-way ANOVA or unpaired t-test.

**Supplemental Figure 8 (Relates to Figure 6).** Locomotor activity does not correlate with vSUB-aBNST in vivo calcium activity. (**A**) Schematic of dual virus injection and monitoring in vivo calcium activity of aBNST projecting vSUB cells in control (pre), ARS, and stress recovery periods (post). Locomotor activity before and after ARS in females (**B**; *P= 0.0148, n=7) and males (**D**; **P=0.010, n=7). Locomotor activity does not correlate with in vivo calcium signal in females (**C**; Pre slope P=0.0992; Post slope P=0.3153, n=7) or males (**E**; Pre slope P=0.9324; Post slope P=0.6563, n=7). Numbers in the legend represent the numbers of cells/animals. Statistical significance was determined by a paired t-test or simple linear regression.

**Supplemental Figure 9 (Relates to Figure 7).** Pharmacologic manipulation of sympathetic or corticosterone signaling does not impact PV-vSUB-RS inhibition in females. (**A**) A Cre-dependent ChIEF AVV was injected into the vSUB of PV-Cre female mice and optogenetically evoked IPSCs from PV interneurons were recorded from RS cells 24-hr after saline, sympathetic block (symp), or corticosterone block (cort) pretreatment. (**B**) Input-output summary graph with representative traces (left) and slope (right) for IPSCs recorded in RS cells after drug controls (**B**; *P*=0.5048; slope, *P*=0.4583; saline n=14/3, symp n=16/3, cort n=14/3). (**C**) Strontium-mediated aIPSC amplitudes after optogenetic stimulation for RS cells (**C**; P= 0.2251; saline n=13/3, symp n=13/3, cort n=13/3). (**D**) PPR (50ms) measurements from RS cells (**D**; *P*=0.2865; saline n=14/3, symp n=16/3, cort n=14/3). (**E**,**I**) A Cre-dependent ChIEF AVV was injected into the vSUB of PV-Cre female mice, mice received a symp (**E**) or cort (**I**) pretreatment before ARS or underwent control conditions, and optogenetically evoked IPSCs from PV interneurons were recorded from RS cells 24-hr later. (**F,J**) Input-output summary graph with representative traces (left) and slope (right) for IPSCs in RS cells in symp (**E**; *P*=0.9889; slope, *P*=0.9140; control n=12/3, symp=15/4) or cort (**i**; *P*=0. 8122; slope, *P*=0.7611; control n=12/3, cort n=13/3) studies. (**G**,**K**) Strontium-mediated aIPSC amplitudes from RS cells after symp (**G**; *P*=0.1952; control n=12/3, stress n=16/4) or cort (**K**; *P*=0.0612, control n=13/3, cort n=13/3) studies. (**H**,**L**) PPR (50ms) measurements in RS cells in symp (**H**; *P*=0.4786, control n=12/3, symp n=15/3) or cort (**L**; *P*=0.3405, control n=12/3, cort n=13/3) studies. Numbers in the legend represent the numbers of cells/animals. Statistical significance was determined by a 1-way ANOVA, 2-way ANOVA, or unpaired t-test.

**Supplemental Figure 10 (Relates to Figure 8).** Pharmacologic manipulation of sympathetic or corticosterone signaling does not impact PV-vSUB-RS inhibition in males. (**A**) A Cre-dependent ChIEF AVV was injected into the vSUB of PV-Cre male mice and optogenetically evoked IPSCs from PV interneurons were recorded from RS cells 24-hr after saline, sympathetic block (symp), or corticosterone block (cort) pretreatment. (**B**) Input-output summary graph with representative traces (left) and slope (right) for IPSCs recorded in RS cells after drug controls (**B**; *P*=0.6765; slope, *P*=0.2248; saline n=12/3, symp n=13/3, cort n=12/3). (**C**) Strontium-mediated aIPSC amplitudes after optogenetic stimulation for BS cells (**C**; P= 0.2585; saline n=11/3, symp n=13/3, cort n=12/3). (**D**) PPR (50ms) measurements from RS cells (**D**; *P*=0.9248; saline n=12/3, symp n=13/3, cort n=12/3). (**E**,**I**) A Cre-dependent ChIEF AVV was injected into the vSUB of PV-Cre female mice, mice received a symp (**E**) or cort (**I**) pretreatment before ARS or underwent control conditions, and optogenetically evoked IPSCs from PV interneurons were recorded from RS cells 24-hr later. (**F,J**) Input-output summary graph with representative traces (left) and slope (right) for IPSCs in RS cells in symp (**E**; *P*=0.6989; slope, *P*=0.3676; control n=14/3, symp=11/3) or cort (**I**; *P*=0.8999; slope, *P*=0.6435 ; control n=14/3, cort n=18/3) studies. (**G**,**K**) Strontium-mediated aIPSC amplitudes from RS cells after symp (**G**; *P*=0.4261; control n=12/3, stress n=12/3) or cort (**K**; *P*=0.9314, control n=13/3, cort n=14/3) studies. (**H**,**L**) PPR (50ms) measurements in RS cells in symp (**H**; *P*=0.2628, control n=11/3, symp n=14/3) or cort (**L**; *P*=0.8935, control n=14/3, cort n=18/3) studies. Numbers in the legend represent the numbers of cells/animals. Statistical significance was determined by a 1-way ANOVA, 2-way ANOVA, or unpaired t-test.

